# A Network Approach to Identify Biomarkers of Differential Chemotherapy Response Using Patient-Derived Xenografts of Triple-Negative Breast Cancer

**DOI:** 10.1101/2021.08.20.457116

**Authors:** Varduhi Petrosyan, Lacey E. Dobrolecki, Lillian Thistlethwaite, Alaina N. Lewis, Christina Sallas, Ramakrishnan Rajaram, Jonathan T. Lei, Matthew J. Ellis, C. Kent Osborne, Mothaffar F. Rimawi, Anne Pavlick, Maryam Nemati Shafaee, Heidi Dowst, Alexander B. Saltzman, Anna Malovannaya, Elisabetta Marangoni, Alana L.Welm, Bryan E. Welm, Shunqiang Li, Gerburg Wulf, Olmo Sonzogni, Susan G. Hilsenbeck, Aleksandar Milosavljevic, Michael T. Lewis

**Author notes:** Senior coauthors (Aleksandar Milosavljevic, Michael T. Lewis).

## Abstract

Triple negative breast cancer (TNBC) is a highly heterogeneous set of diseases that has, until recently, lacked any FDA-approved, molecularly targeted therapeutics. Thus, systemic chemotherapy regimens remain the standard of care for many. Unfortunately, even combination chemotherapy is ineffective for many TNBC patients, and side-effects can be severe or lethal. Identification of predictive biomarkers for chemotherapy response would allow for the prospective selection of responsive patients, thereby maximizing efficacy and minimizing unwanted toxicities. Here, we leverage a cohort of TNBC PDX models with responses to single-agent docetaxel or carboplatin to identify biomarkers predictive for differential response to these two drugs. To demonstrate their ability to function as a preclinical cohort, PDX were molecularly characterized using whole-exome DNA sequencing, RNAseq transcriptomics, and mass spectrometry-based total proteomics to show proteogenomic consistency with TCGA and CPTAC clinical samples. Focusing first on the transcriptome, we describe a network-based computational approach to identify candidate epithelial and stromal biomarkers of response to carboplatin (*MSI1, TMSB15A, ARHGDIB, GGT1, SV2A, SEC14L2, SERPINI1, ADAMTS20, DGKQ*) and docetaxel (*ITGA7, MAGED4, CERS1, ST8SIA2, KIF24, PARPBP)*. Biomarker panels are predictive in PDX expression datasets (RNAseq and Affymetrix) for both taxane (docetaxel or paclitaxel) and platinum-based (carboplatin or cisplatin) response, thereby demonstrating both cross expression platform and cross drug class robustness. Biomarker panels were also predictive in clinical datasets with response to cisplatin or paclitaxel, thus demonstrating translational potential of PDX-based preclinical trials. This network-based approach is highly adaptable and can be used to evaluate biomarkers of response to other agents.

## Introduction

Triple-negative breast cancer (TNBC) is an aggressive clinical subtype of breast cancer that is characterized by the absence of expression of the steroid hormone receptors (HR) estrogen receptor alpha (ESR1) and progesterone receptor (PGR), as well as the absence of overexpression and/or genomic amplification of oncogenic epidermal growth factor receptor 2 (ERBB2 or HER2)(Mehanna et al., 2019). Relative to patients diagnosed with HR+ and ERBB2-driven breast cancers, patients with TNBC are typically diagnosed at a younger age, are at a higher risk of relapse, and have lower overall survival rates. The median overall survival rate for patients with metastatic TNBC remains less than 18 months (Garrido-Castro et al., 2019). Unlike HR+ and ERBB2-driven tumors, which are treated with either endocrine therapy or ERBB2-targeted therapies, respectively, TNBC largely lacks biomarker-guided selection for treatment with targeted agents. Thus, treatment options for many TNBC patients remain limited to regimens using cytotoxic chemotherapies, which also lack predictive biomarkers for guided agent selection.

In recent years, there have been some breakthrough advances for treating TNBC more effectively. Immune checkpoint inhibitors including Atezolizumab, which targets Programmed Cell Death Ligand 1 (PD-L1), and Pembrolizumab, which targets Programmed Cell Death Receptor 1 (PD-1), are approved by the Food and Drug Administration (FDA) in locally advanced, non-resectable and metastatic patients. Atezolizumab is given with nab-paclitaxel, and Pembrolizumab is given with a broader range of chemotherapeutics. Pembrolizumab has also been approved with chemotherapy (various) in high risk primary TNBC in the neoadjuvant setting. In addition to immunotherapies, PARP inhibitors olaparib and talazoparib are also approved for HER2-negative patients with advanced disease that carry a germline BRCA1/2 mutation, and who have had prior chemotherapy. Finally, the FDA recently approved sacituzumab govitican, for locally advanced, unresectable primary tumors, as well as metastatic TNBC. Sacituzumab govitican acts by recognizing Trop2 (trophoblast cell-surface antigen 2) with an antibody conjugated to SN38 (the active metabolite of irinotecan targeting topoisomerase I). Patients treated with this therapy must have received two prior systemic treatments, with at least one of these being for metastatic breast disease (i.e. chemotherapy). Strikingly, despite these major advances, systemic chemotherapies remain and integral part of the TNBC treatment landscape.

Reduction in tumor volume is relevant in multiple clinical settings. In the neoadjuvant setting, systemic treatment is used to shrink tumors prior to surgery. There are two main reasons that tumor responsiveness is important in this clinical setting. First, patients whose tumors achieve a pathologic complete response (pCR) show better overall and disease-free survival (Spring et al., 2020). Second, even in those patients not showing pCR, tumor shrinkage can enhance surgical outcomes. In the adjuvant setting, systemic treatment is used in an attempt to eliminate residual tumor cells after surgery that may be capable of re-growing a tumor either as a local or distant recurrence. Finally, in the metastatic setting, agents are used in an attempt to either eliminate tumors, or reduce tumor burden, to extend life. Thus, development of methods to determine which patients, in which clinical setting, will respond to which agents, systemic chemotherapies or otherwise, is an important clinical/translational goal.

With respect to single agent/regimen treatment response in the neoadjuvant setting (taxane, platinum, anthracyclin/cytoxan), only 25-33% of TNBC patients achieve pCR to any given treatment (Caparica et al., 2019). This mediocre response rate has led to the combination of chemotherapeutic agents, with the combinations and dose schedules determined to be most effective identified over time in human clinical trials. Combination chemotherapy is now first line standard of care for TNBC in the neoadjuvant setting, with patients receiving up to five chemotherapeutic agents over the course of treatment (Wahba and El-Hadaad, 2015). Even with the most recently used chemotherapy combinations, pCR rates still only reach 55-65% (Loibl et al., 2018; Minckwitz et al., 2014; Sharma et al., 2017; Sikov et al., 2014). Because 35-45% of patients do not respond even when their tumors are challenged with multiple cytotoxic chemotherapy agents, it is clear that many patients receive toxic, and ultimately ineffective, treatments for little or no clinical benefit.

If systemic combination chemotherapy is to remain at the forefront of treatment for TNBC, in any clinical setting, it is critical that molecular biomarkers of differential response to individual chemotherapy agents be identified. If successful, patients can be selected prospectively as a likely responder to one or more agents, and then treated with the agent(s) most likely to be effective against their unique tumor. If successful, efficacy rates could increase dramatically. Further, there should be the opportunity for dose de-escalation, which should decrease the frequency of severe, or life-threatening toxicities. Finally, these data may also provide insights into molecular mechanisms of chemotherapy resistance that may be targetable to enhance responses.

Historically, attempts to develop response predictors have been limited to either *in vitro* cell line-based studies, cell line xenografts, or human clinical trials. However, to date, predictive molecular signatures of chemotherapy response are not used clinically. More recently, patient-derived xenograft (PDX) models of human breast cancer have emerged as potential surrogates for their tumor of origin. We and others have shown remarkable biological consistency between patient tumors and their corresponding PDX with respect to histology, cellular heterogeneity, biomarker expression, mutations, genomic copy number alterations, variant allele frequencies, and mRNA expression patterns (Dobrolecki et al., 2016; Echeverria et al., 2018; Evrard et al., 2019; Powell et al., 2020; Savage et al., 2020; Whittle et al., 2015; Woo et al., 2019; Zhang et al., 2013a). Most importantly, we and others have made progress in demonstrating that treatment responses in PDX are qualitatively similar to those of the tumor-of-origin (Savage et al., 2020; Whittle et al., 2015; Zhang et al., 2013a). Based on these commonalities, and the availability of a comparatively large collection of TNBC PDX, we hypothesized that collections of PDX models can be used as a cohort analogous with human cohorts in clinical trials. If so, it should be possible to derive pre-clinically and clinically relevant molecular signatures of treatment response using a collection of PDX as the discovery platform.

To evaluate the degree to which our PDX collection could function as a “patient cohort” in preclinical trials that might also inform tumor responses in clinical trials, we conducted a baseline proteogenomic characterization using DNA whole exome sequencing (copy number alterations and mutations), mRNA transcriptomics (deep RNAseq), and mass spectrometry-based total proteomics) and compared these data with large patient datasets to demonstrate the range of representation of the PDX models. We then aggregated single agent responses to docetaxel or carboplatin across three recently completed PDX-based preclinical trials designed to approximate human equivalent dosing and scheduling in the mouse (to be described in full elsewhere) for analysis. We ultimately selected mRNA from other -omic data types for generation of differential response signatures as it was the deepest omics signature, and we could identify both PDX-based and clinical datasets with which to validate our observations.

Standard methods of identifying genes informative for prediction of response and resistance include simple correlation of gene expression with the phenotype of interest, recursive feature elimination in machine-learning models, and the identification of hub genes with graph-based methods such as WGCNA (Weighted Gene Co-expression Network Analysis) (Jia et al., 2020; Tadist et al., 2019; Tang et al., 2018; Zhao et al., 2010). Determining the correlation between genes and the phenotype of interest is perhaps the most straightforward approach for the identification of potentially informative genes. A problem with using gene-phenotype correlations is that a gene may be correlated to response not because of a direct role in response, but because it is correlated with other factors that are functionally relevant for response. Another standard approach is to identify genes that add predictive power to a machine learning model of response. An example of this approach is the recursive feature elimination method. This method begins with all potentially informative features and eliminates features recursively based on the importance of individual features to the model. Unfortunately, recursive feature elimination can eliminate features which, while weak on their own, could contribute predictive power in the context of other features. Network-based methods such as WGCNA have also been developed to identify informative and biologically connected sets of genes. These methods are often more reliable for the identification of biomarkers than single gene comparison methods. WGCNA is a widely exploited network method in the field, and it has been used to identify informative modules and their hub genes in complex diseases including cancer and Alzheimer’s (Di et al., 2019; Du et al., 2020; Giulietti et al., 2017; Huang et al., 2020; Liao et al., 2020; Liu et al., 2017; Qiu et al., 2019; Zhang et al., 2018; Zhao et al., 2010).

The WGCNA approach is not without limitations. The standard application of WGCNA includes building a network over all samples, detecting informative modules through the correlation of the representative module eigengene profile to the phenotype of interest, and finally the identification of highly connected hub genes in informative modules. There are three primary issues with the standard WGCNA approach. The modules identified by WGCNA can be large and thus biologically unwieldy. Results may also be associated with inter-sample biological heterogeneity rather than to actual response. Furthermore, the use of module eigengenes to determine if a module is associated with response may miss more highly informative genes in an otherwise less informative module. This happens when a large module (hundreds of genes) contains a small, more highly informative, submodule whose signal is diluted by the presence of the large number of less informative genes.

To test our hypothesis that PDX-based cohorts can yield translationally relevant predictive information, and to address one of the limitations of WGCNA, we adapted a newly developed network algorithm, CTD (Thistlethwaite et al., 2021), to “Connect The Dots” in our molecular data relative to single agent chemotherapy responses. Here, CTD is used in addition to WGCNA to identify highly connected sets of analyte nodes, and to assign an upper-bounded p-value to sets of nodes. While other network-based algorithms also estimate p-values, they use permutation testing which is not feasible for large gene co-expression graphs with thousands of genes. Our ability to assign p-values without the need for permutation testing allows the identification of small sets of nodes that are significantly associated with a phenotype. This algorithmic advancement allowed for the identification of a specific set of genes that were differentially associated with docetaxel and carboplatin response/resistance in our PDX cohort. These PDX-derived informative panels were validated for their predictive power in other PDX collections, as well as in two clinical datasets. If validated in subsequent prospective clinical trials, this approach to feature selection coupled with PDX-based discovery could accelerate development of predictive signatures that are useful clinically.

## Results

### PDX models of TNBC closely resemble human tumors represented in The Cancer Genome Atlas (TCGA) and Clinical Proteomic Tumor Analysis Consortium (CPTAC) Cohorts

At BCM, we have amassed a collection of 169 PDX models of human breast cancer that is now large enough that a subset of the collection could potentially be used as a pre-clinical cohort whose biological characteristics should closely resemble clinical cohorts (publicly available models can be evaluated on https://pdxportal.research.bcm.edu/pdxportal). These models are summarized in *Supplemental Figure 1* which was generated with ComplexHeatmap (Gu et al., 2016).

For this study, we make use of a subset of 50 PDX models of TNBC, all of which were treated with single agent docetaxel or carboplatin at human equivalent doses, and in a preclinical platform that closely mimics a multicycle clinical trial in human patients(Izumchenko et al., 2017). This PDX collection shows a wide range of responses to docetaxel and carboplatin, with some PDX being cross-resistant or cross-responsive, but others showing differential responses to these two agents, suggesting that molecular signatures of differential treatment response could be generated that might also have predictive power in clinical datasets (Zhang et al., 2013b).

To evaluate the quality of our -omic data, and to determine if our collection of TNBC PDX reflected a significant cross section of TNBC clinical samples represented in TCGA (genome and transcriptome) and CPTAC (proteome) to be considered a representative preclinical cohort, we compared DNA, RNA, and protein-based assays.

At the DNA level, we first compared genomic copy number alteration patterns in our cohort relative to TCGA samples using WES data. Figure 1A shows that the overall pattern of gains and loses across our PDX cohort and the TCGA cohort are qualitatively similar upon visual inspection. Similar results were obtained for ER+ and HER2+ PDX, but our sample size was not large enough for full analysis (*Supplemental Figure 2A and 2B*). To demonstrate similarities in a more quantitative manner, we clustered the gain and loss patterns of both the human and PDX cohorts. Figure 1B shows these hierarchical clustering results. Not only could PDX gain/loss patterns be compared directly to TCGA samples, but the results also show that our PDX collection represents a wide range of TNBC at the copy number alteration level.

**Figure 1:**
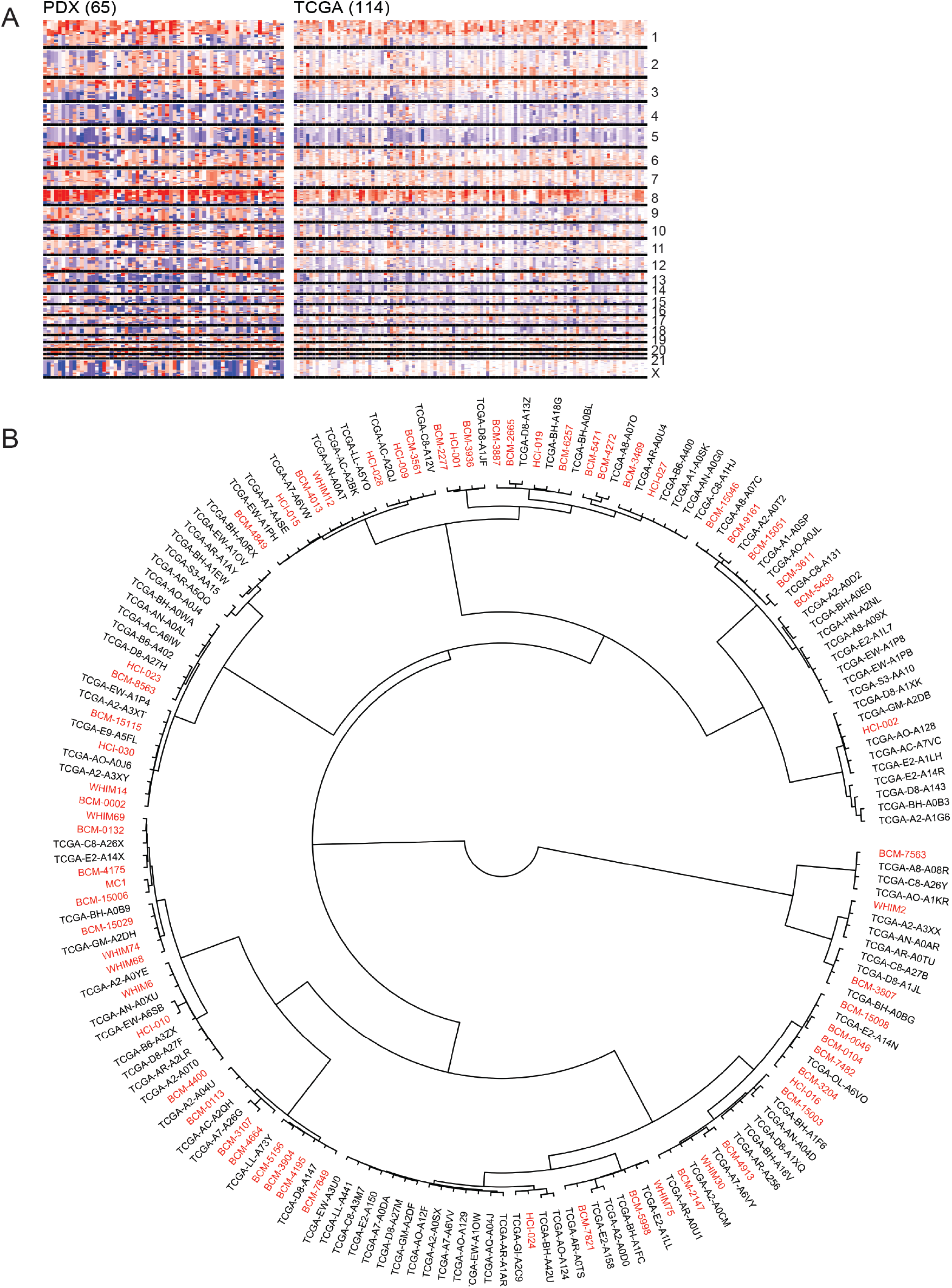
Copy number variation is quantitatively and qualitatively similar in TNBC PDX models and TNBC clinical samples. Figure 1A: Copy number variation comparison between the PDXs and the TNBC clinical samples in TCGA with a heatmap. Figure 1B: Hierarchical clustering of PDXs among the TCGA TNBC samples with copy number variations to illustrate the distribution of PDX models within human clinical samples. PDX samples are in red and TCGA samples are in black.

Next, we evaluated the spectrum of mutations in our TNBC PDX cohort relative to TNBC in TCGA to determine whether the distribution of mutations was indeed similar. Figure 2A demonstrates that the spectrum of genes most highly mutated in TCGA are also mutated at largely the same frequencies in PDX. Similar results were observed for ER+ and HER2+ PDX models, (*Supplemental Figure 3A and 3B*, respectively). While our sample size for either the ER+ or HER2+ subtypes were not large enough to be representative, these results suggested that the mutation spectrum observed in our PDX collection is representative of the mutation spectrum observed in human patients, particularly TNBC patients.

**Figure 2:**
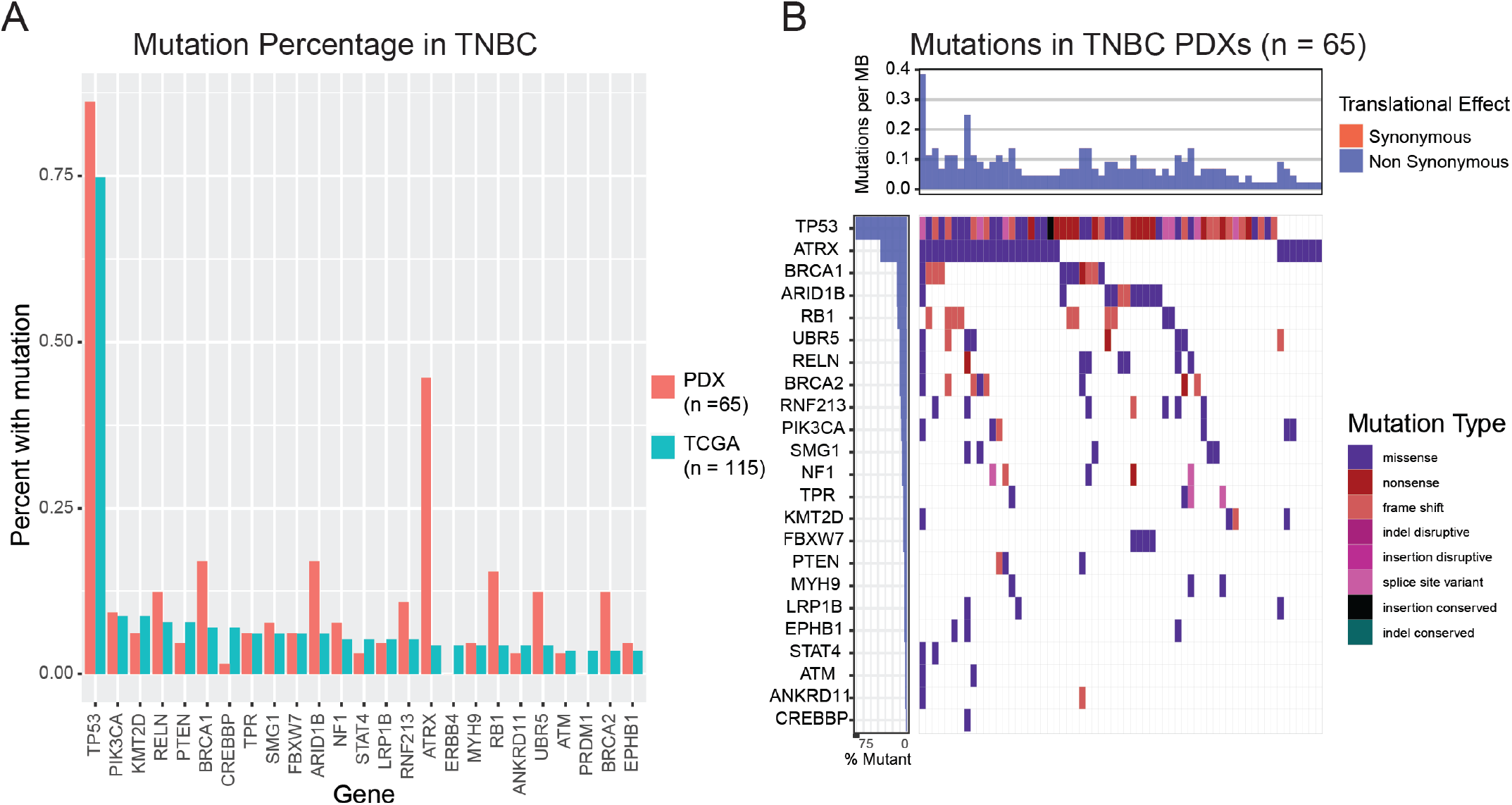
Mutational load is similar in TNBC PDXs and TNBC clinical samples. Figure 2A: Mutational load of PDX models and TNBC TCGA samples show similar mutational profiles in highly mutated genes. Figure 2B: Waterfall plot showing the distribution of the types of mutations found in the PDX models.

With respect to gene expression analysis at the transcriptomic level, deep RNAseq (~200M reads per sample) GSEXXXXXX data from untreated PDX models were hierarchically clustered with patient tumors represented in TCGA using the PAM50 signature (*Figure 3A*) (Koboldt et al., 2012). Importantly, the PDX models do not cluster together as a single group. Rather, PDX were interspersed among TCGA samples, with basal-like PDX being distributed into multiple groups across the full range observed in TCGA.

**Figure 3:**
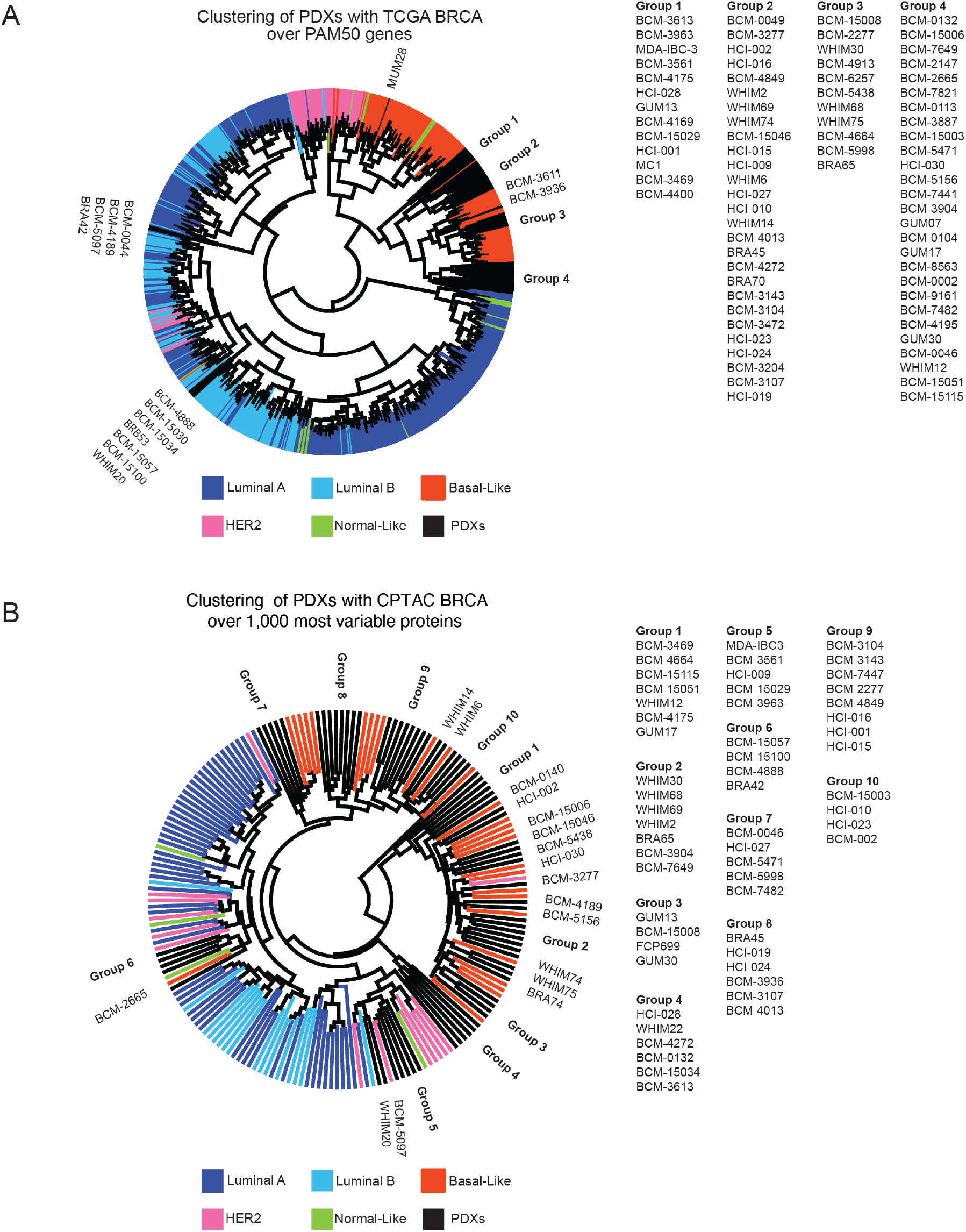
Deep RNAseq and proteomics reveal similarities between PDX models and TCGA clinical breast cancer samples. Figure 3A: Hierarchical clustering of TCGA and PDX models over the PAM50 gene signature. Figure 3B: Hierarchical clustering of TCGA and PDX models over the top 1,000 most variable proteins.

With respect to gene expression analysis at the proteomic level, we detected 8346 human proteins in at least half of our models by mass spectrometry. As with the RNAseq analysis, PDX models were again distributed across the samples represented in the CPTAC proteomic dataset using the top 1000 most variable proteins (*Figure 3B*). The clustering pattern was remarkably like that derived from mRNA data, again with two main groups clustering with luminal B tumors, and the majority clustering in the basal-like group, with PDX models distributed across the full spectrum of basal-like tumors.

### Responses Of PDX Models To Human-Equivalent Doses Of Docetaxel Are Largely Consistent With Clinical Responses Of The Tumor-Of-Origin When Challenged With A Taxane

To evaluate concordance of treatment responses across a range of chemotherapies, we compared the “best clinical response” of PDX using our modified RECIST 1.1 classification to clinical and/or pathologic responses in their tumor of origin when challenged with the same, or same class, of agent. Results are shown in Tables 1-3.

In the patient matched PDX models, treatment with 20mg/kg docetaxel (Table 1), 14/30 (47%) showed unqualified concordance with the tumor of origin. An additional 11 (37%) showed “qualified” concordance, in which the PDX response fell into an adjacent RECIST response category in the patient. Given that the preclinical regimen we established may not yet be fully equivalent to that used in a human clinical trial, we reasoned that the use of qualified concordance was justified. If this argument is accepted, a remarkable 84% of observations were considered concordant. Only 4/30 PDXs received an unqualified discordant response from the original tumor ((“No”) shown here in pink).

For the four unqualified non-concordant PDX models (BCM-3611, BCM-4913, BCM-7482, and HCI-015) three tumors of origin showed Progressive Disease (PD), whereas the corresponding PDX (BCM-3611, BCM-4913, and HCI-015) showed a Partial Response (PR). The reason for the enhanced response relative to the tumor-of-origin is not known. The remaining PDX, BCM-7482, showed PD while the tumor of origin showed a PR. For this line, we hypothesized that the difference in response may be because we evaluated docetaxel at the low human equivalent dose (20mg/kg) rather than the high human equivalent dose (30mg/kg) to which the tumor of origin would likely have been exposed. If so, BCM-7482 may show response to the higher dose, and thus then show concordance. To test this possibility, we challenged BCM-7482 with four cycles of docetaxel at the higher dose. Several other lines were also evaluated to see if response to 30mg/kg docetaxel enhanced response beyond that of 20mg/kg (HCI-015 was not evaluated). As hypothesized, BCM-7482 then became concordant. Thus, overall, only 10% (3/30) of the PDX evaluated showed unqualified discordance.

### Concordance Of Response With The Tumor-of-origin: PDX Responses To Carboplatin Or AC Were Equivocal

With respect to platinum-based agents, we identified five PDX-matched patients who had been treated with a platinum-containing regimen, none of which were single agent. Of these, there were two unqualified concordances and one qualified concordance (most likely due to the addition of radiation with cisplatin), and two unqualified discordances (Table 2). In the two discordant cases, the patients were treated with combination carboplatin/taxane, which might complicate the interpretation of the response results. Thus, it remains to be determined whether PDX responses to platinum-containing regimens recapitulate patient tumor-of-origin responses.

Finally, with respect to multi-cycle AC (anthracycline and cyclophosphamide), nine patient-matched PDX could be evaluated for concordance. In contrast to docetaxel-treated PDX, AC-treated PDX showed only three unqualified concordances and two qualified concordances. Four of nine (44%) showed unqualified discordance compared to the tumor of origin. In most cases, the clinical response of the tumor of origin was PR, whereas the corresponding PDX showed PD. This is most likely due to the fact that we could not achieve human equivalent doses for either the doxorubicin or cyclophosphamide. Thus, PDX responses to AC are unlikely to be fully reflective of the responses in patients, at least under the conditions achievable in SCID/Bg mice.

### Identification Of Multigene Biomarker Panels Associated With Differential Chemotherapy Response Using a WGCNA Only Vs. a Combined WGCNA/CTD Approach

Based on the conservation of biology in the -omics datasets, the high degree of concordance of responses in docetaxel-treated PDX/patient tumor pairs, and the possibility that PDX may ultimately recapitulate responses to carboplatin, we chose to focus our effort on taxane and platinum differential response prediction.

Because the RNAseq transcriptomic data were the most robust, and could be validated in other datasets, we chose to focus on these data first. For each PDX, epithelial gene expression and stromal gene expression was evaluated by separating the RNAseq reads by species using Xenome. Separation allowed us to evaluate both epithelial and stromal genes that may be associated with resistance for each approach. For both the WGCNA and combined WGCNA/CTD approach, we first built two networks for each cell type (stromal or epithelial)/response combination using WCGNA only (biweight midcorrelation).

One of these graphs was built over all the PDX models, while the other was built just over the responsive PDXs (CR and PR). Because gene-gene correlation could be caused by heterogeneity in the samples that is not related to the responsiveness of the samples to the agents in question, we then pruned, or removed edges, that were found in the graph built over just the responsive PDXs to ensure that we eliminated noise not related to response. By removing these edges, we both eliminated gene-gene variance that may not be associated with response and made the networks more sparse, and thus easier to evaluate. This approach produced four pruned networks that were specific to both a tissue compartment and chemotherapy treatment (i.e., human epithelial carboplatin, human epithelial docetaxel, murine stromal carboplatin, murine stromal docetaxel).

Pruned networks were then broken into modules with WGCNA using the standard approach. The method identified between 3 and 99 gene modules for each of the graphs. These modules were large (30-200 genes each), and not biologically tractable as they contained too many genes representing several pathways, thus not allowing us to determine which specific genes were most closely related to response in a straightforward manner.

As mentioned in the introduction, the WGCNA approach may overlook modules with informative genes whose signal is overwhelmed by less-informative genes. After the identification of informative modules, WGCNA identifies hub genes, but these the hub genes are not necessarily the genes that carry the predictive information. To address these issues, we applied CTD to determine which sets of genes in these large WGCNA modules were significantly connected in the pruned networks.

CTD is an information theoretic-based network method that identifies patterns of connectedness between analytes and assigns p-values to highly connected subsets of analytes within large modules without computationally-costly permutation testing (*Figure 4A*) (Thistlethwaite et al., 2021). With respect to gene expression data, CTD is used to “connect the dots” and identify these highly connected gene sets by finding subsets that are more connected in a weighted graph than by chance. Because response has not been added as a node in the expression graph, we hypothesize that significantly connected sets within the large modules may be connected due to their latent connection to response (*Figure 4B*). For each of the pruned networks, we identified small CTD submodules that contained fewer genes than the number of PDXs in our analysis (n=45). This then allowed for the generation of generalized linear models (GLMs), and the identification of specific genes that are predictive for response to the two agent classes chosen. The p-value calculated by CTD for these modules, as well as the significance of their linear regression with response, and whether the larger module was identified as informative by WGCNA can be found in *Supplemental Table 1*.

**Figure 4:**
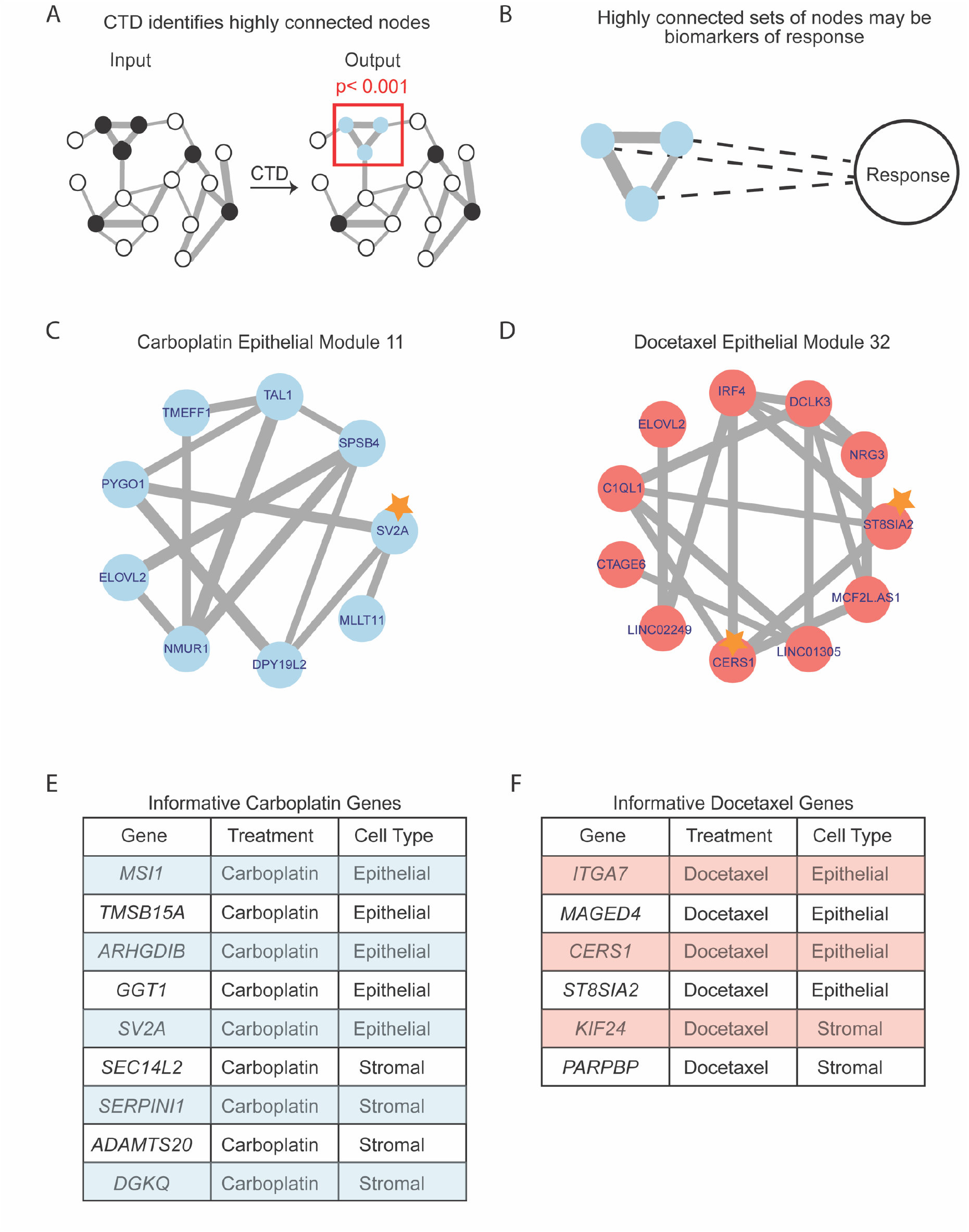
CTD connects the dots to identify multigene biomarker panels of response to carboplatin and docetaxel. Figure 4A: CTD is a network method that can be used to “connect the dots” and identify highly connected sets of genes. CTD takes a weighted graph and a set of nodes of interest (grey) and outputs a set of highly connected genes (purple) with a p-value of connectedness. Figure 4B: Highly connected nodes may be connected through their latent connection to response. Figure 4C: CTD carboplatin epithelial Submodule 5 which contains 3 informative genes highlighted with a star. Figure 4D: CTD docetaxel epithelial Submodule 23 which contains one informative gene highlighted with a star. Figure 4E: Table of all the informative genes for carboplatin identified with the WGCNA/CTD approach. Figure 4F: Table of all the informative genes for docetaxel identified with the WGCNA/CTD approach.

An example of an informative submodule for the carboplatin epithelial network is shown in *Figure 4C*, while an example of an informative docetaxel epithelial submodule is shown in *Figure 4D*. For carboplatin, five genes were identified as informative for response in the epithelial network (*MSI1, TMSB15A, ARHGDIB, GGT1, SV2A*) and four genes were identified as informative in the stromal network (*SEC14L2, SERPINI1, ADAMTS20, DGKQ*) (*Figure 4E)*. For docetaxel, four genes were identified as informative in the epithelial network (*ITGA7, MAGED4, CERS1*, *ST8SIA2*) and two genes were identified as informative in the stromal network (*KIF24, PARPBP*) (*Figure 4F)*.

We then combined the stromal and epithelial RNAseq expression profiles for each of the PDX tumor models to generate a pseudo-bulk expression profile for each PDX. To show the directionality of these informative genes relative to response, we plotted the expression of our informative panel in the pseudo-bulk expression profiles for the carboplatin genes (*Figure 5A)* and the docetaxel genes (*Figure 5B*) relative to quantitative drug responses. In both cases, an expression gradient can be observed across PDX with respect to response.

**Figure 5:**
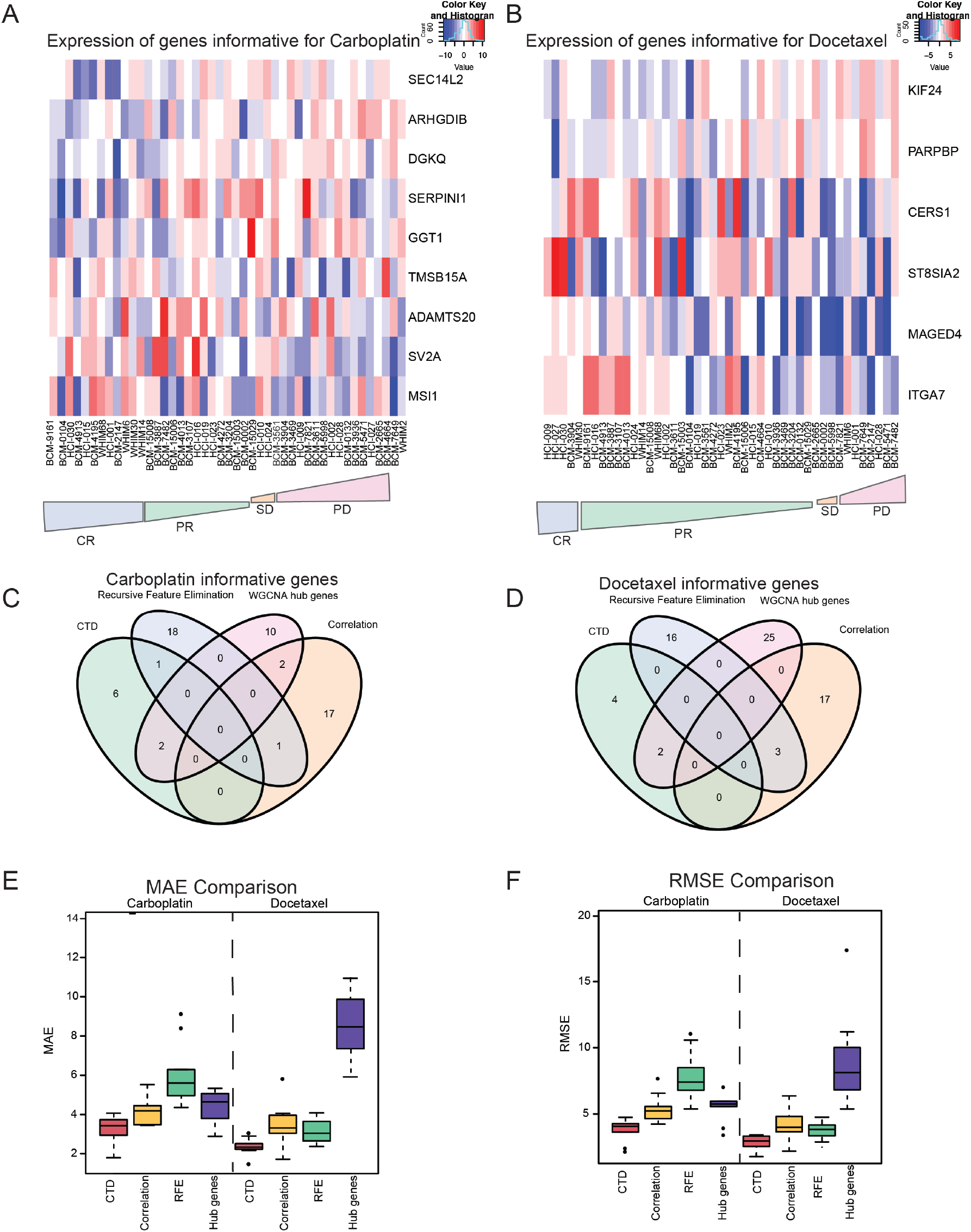
CTD outperforms other feature selection methods when selecting informative genes and is predictive in our PDX cohort. Figure 5A: Heatmap of informative genes for carboplatin across all PDXs from most responsive (left) to most resistant (right). CR – complete response, PR-partial response, SD – stable disease, PD – progressive disease Figure 5B: Heatmap of informative genes for docetaxel across all PDXs from most responsive (left) to most resistant (right). CR – complete response, PR-partial response, SD – stable disease, PD – progressive disease Figure 5C: Overlap between genes predicted to be informative for carboplatin. Figure 5D: Overlap between genes predicted to be informative for docetaxel. Figure 5E: MAE comparison of 4 different methods (red = WGCNA/CTD approach, orange = correlation, green = recursive feature extraction, purple = using the WGCNA hub genes) Figure 5F: RMSE comparison of 4 different methods (red = WGCNA/CTD approach, orange = correlation, green = recursive feature extraction, purple = using the WGCNA hub genes)

### Protein Multigene Biomarker Evaluation

Because our total proteome data were more sparse, WGCNA did not perform well enough to evaluate the WGCNA/CTD approach. To determine if the multigene biomarkers that were identified with RNA expression data were also predictive of response using protein data, we leveraged a mass spectrometry-based total proteomics dataset for the same PDX models. The proteins were separated into those expressed in cancer cells and those expressed in the murine stroma by the gpGrouper algorithm (Saltzman et al., 2018). We then identified proteins whose expression was significantly correlated to response and compared the list of identified proteins to both our multigene markers and the informative submodules *(Supplemental Table 2).*

For carboplatin, we found that three of our informative genes at the mRNA level were also significantly correlated (p < 0.05) with response at the protein level (*MSI1, ARHGDIB, DGKQ,*), as well as two other genes (*MAGED4, MAGED4B*) that were found in gene expression submodules that were informative for carboplatin response *(Supplemental Table 2).* The Fisher’s exact test for the intersection of informative genes identified by the WGCNA/CTD approach and proteins correlated to response was significant with a p-value of 0.002.

For docetaxel, only one of the proteins significantly correlated with response (KIF18B) was also identified in a CTD submodule. The Fisher’s exact test was marginally significant for this comparison with a p-value of 0.1. The identification of a subset of informative gene biomarkers through an independent protein analysis further increased our confidence in the multi-gene RNA biomarkers.

### Enrichment By Text Mining Using The WGCNA/CTD Approach And Literature Search

To determine if the WGCNA/CTD method enriched for genes that were previously shown to be associated with chemotherapy resistance, we employed a text mining approach. iTextMine (Ren et al., 2018) was used to query which of the genes identified by WGCNA alone, as well as in our informative panel were also identified in other studies as linked to the term “chemoresistance”. Of the 15 WGCNA/CTD genes, two (13%) were also identified with the text mining approach (Fisher’s test p-value = 0.06), as opposed to only 16 (4%) of the 390 WGCNA genes (Fisher’s test p-value = 0.11). The genes identified by both the WGCNA/CTD approach and text mining were *MSI1* and *ARHGDIB*. As well as identifying a smaller more biologically tractable set of genes, the addition of CTD to the standard WGCNA approach also enriched for genes that have previously been linked to the term “chemoresistance” in the literature.

To determine more rigorously if our multigene biomarkers were associated with cancer progression or response to therapy previously, we then conducted a literature search. For the carboplatin multigene biomarker panel, we found that: *MSI1* was associated with lung cancer malignancy (Lang et al., 2017) and stem cells in breast cancer (Lagadec et al., 2014), *TMSB15A* is predictive of response in TNBC (Darb-Esfahani et al., 2012), *ARHGDIB* is associated with breast cancer progression and is associated with drug resistance (Bozza et al., 2015; Zhang et al., 2005), *GGT1* is linked to breast cancer progression and prognosis (Staudigl et al., 2015), *SERPINI1* may be associated with the survival of cancer cells (Valiente et al., 2014), and ADAMTS20 is associated with the grade of breast cancer (Guo et al., 2018).

For the docetaxel multigene biomarkers, we found that: *ITGA7* is associated with worse prognosis and migration in breast cancer (Bai et al., 2019; Bhandari et al., 2018) and is a stem cell marker in squamous cell carcinoma(Ming et al., 2016), breast cancer patients with an overexpression of *MAGED4* had worse survival (Jia et al., 2019), *CERS1* is associated with cancer cell death in head and neck squamous cell carcinomas (Dany and Ogretmen, 2015), and *PARPBP* has previously been identified in breast cancer as a marker of therapy resistance (Chen et al., 2020).

Additionally, *SV2A* (Bandala et al., 2015) and *KIF24* (Kim et al., 2015) were found to be overexpressed in breast cancer cell line data.

### Comparison Of The WGCNA/CTD Approach With Other Feature Selection Methods

We then investigated how the combined WGCNA/CTD approach compared to three widely used feature selection methods (correlation, recursive feature elimination with a random forest model, and WGCNA hub genes). For evaluation of the WGCNA only approach, we identified modules whose eigengenes were correlated to response, and then determined which genes had the highest gene significance. For the correlation and recursive feature selection approaches, we identified the top 10 genes (as our WGCNA/CTD approach identified 9 genes for carboplatin and 6 genes for docetaxel) that were related to response in both the stromal and epithelial compartments. While a few genes were identified with multiple methods, most potentially informative genes were unique to the feature selection method used (*Figure 5C*, *Figure 5D*). Genes that were identified as potentially informative with multiple methods included: *RFC5, SV2A, GGT1, MAGED4, MAGED4B, MSI1* for carboplatin, and *ASGR1, IQSEC1, LYRM2, MAGED4, ST8SIA2* for docetaxel, with only *MAGED4* represented in both panels.

### WGCNA/CTD Outperforms Other Methods Of Feature Selection With Respect To Predictive Value

To evaluate CTD as an informative feature selection method, we tested each of these feature selection methods for their performance predicting treatment response using “pseudo-bulk” RNAseq data reconstructed from the PDX epithelial and stromal RNAseq gene expression datasets. We built these pseudo-bulk profiles by summing the epithelial and stromal mRNA expression values for common genes to approximate human clinical bulk RNAseq better. To evaluate the performance of each feature selection method with a linear model within our dataset, we compared the Mean Absolute Error (MAE) (*Figure 5E*) and Root Mean Square Error (RMSE) (*Figure 5F*) from the quantitative response predictions using a 70/30 training/test set splitting approach of the source data, which were reshuffled 10 times. While each feature selection method identified a largely unique set of genes, we found that the WGCNA/CTD approach outperformed the others, including WGCNA alone, when predicting quantitative response of PDX models to both carboplatin and docetaxel (*Figures 5E-5F*).

### Preclinical Validation Of Differential Chemotherapy Response Predictors

For potential validation in other PDX datasets and in clinical trial response assessments, we converted our quantitative response assessments to a qualitative modified RECIST 1.1 classification for each PDX. We then determined how predictive our models were for both Complete Response (CR) vs. all else, and for CR and Partial Response (PR) vs. all else using the full 50 PDX dataset.

For this analysis, 70% of the pseudo-bulk RNAseq samples were used to train quantitative GLMs of response which was tested on the 30% test set. The training and test sets were reshuffled 10 times to ensure that our results were not specific to a training/test set. The GLMs were predictive in the test set for both carboplatin (MAE = 3.21, RMSE = 3.8) and docetaxel (MAE = 2.37, RMSE = 2.87). ROC curves were constructed using LOOCV for each of the treatments for two response cutoffs: complete response (CR) vs. all else, and complete response plus partial response (CR/PR) vs. all else. The AUC for each combination of treatment and response cutoffs were calculated and are as follows: carboplatin CR/PR AUC = 0.80, docetaxel CR/PR = 0.80, carboplatin CR AUC = 0.85, docetaxel CR AUC = 0.64 (*Figure 6A*, *Figure 6B*). Within our PDX dataset, we found that both the carboplatin biomarker panel and the docetaxel biomarker panel were informative for both quantitative and qualitative response.

**Figure 6:**
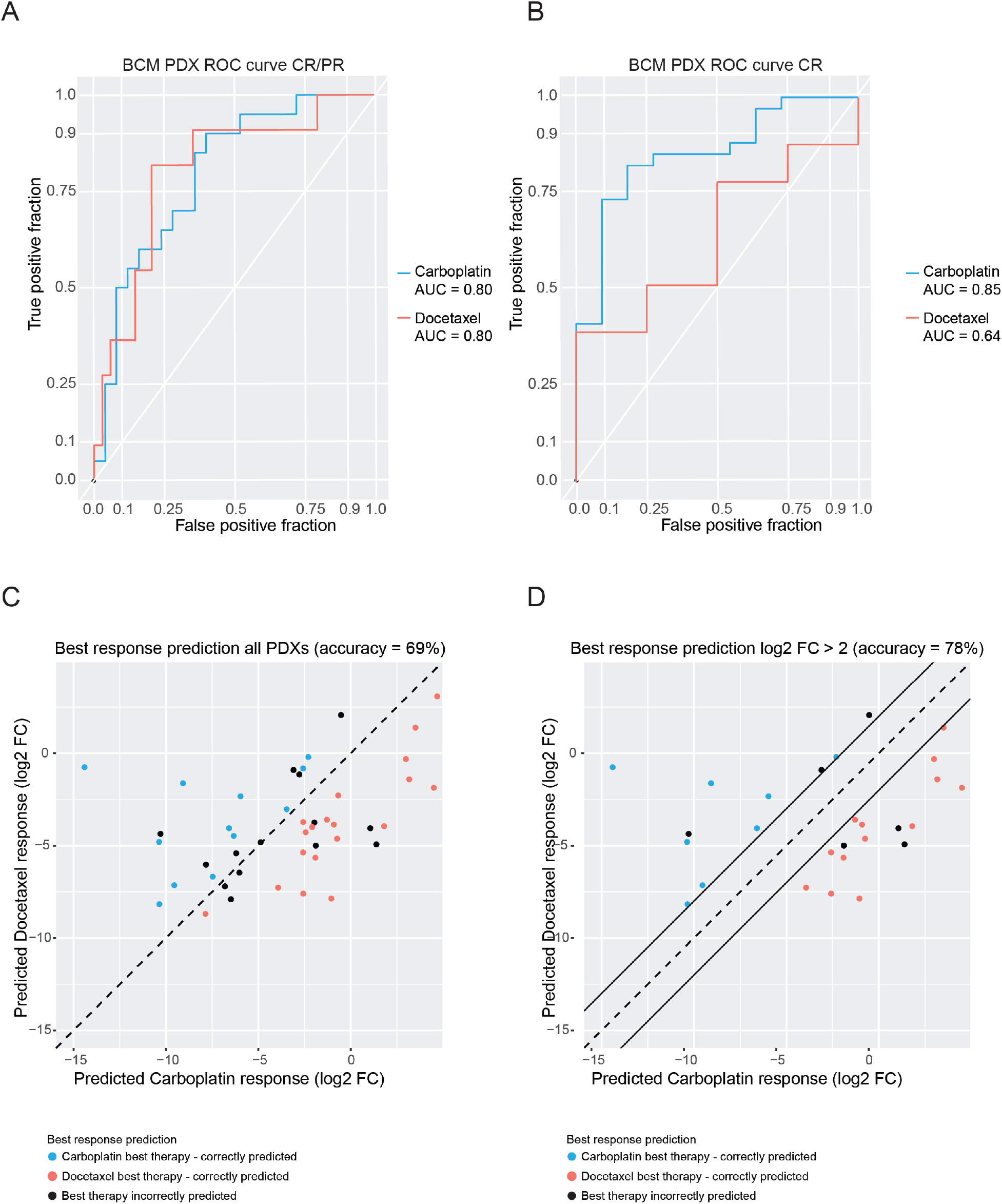
Multigene biomarkers are predictive of qualitative response and are also predictive of the best therapy for each PDX. Figure 6A: ROC for complete and partial response (CR/PR) vs. all else. The ROC for docetaxel is red and the ROC for carboplatin is blue. Figure 6B: ROC for complete partial response (CR) vs. all else. The ROC curve for docetaxel is red and the ROC for carboplatin is blue. Figure 6C: The best response predictions for all PDXs. If both the actual best therapy and the predicted best therapy is carboplatin, the dot is blue. If both the actual best therapy and the predicted best therapy is docetaxel, the dot is red. If the best therapy and the predicted best therapy don’t match the dot is black. Figure 6D: The best response predictions for PDXs that have a predicted log2 fold change of more than 2. If both the actual best therapy and the predicted best therapy is carboplatin, the dot is blue. If both the actual best therapy and the predicted best therapy is docetaxel, the dot is red. If the best therapy and the predicted best therapy don’t match the dot is black.

To be maximally useful, biomarker panels predictive of differential drug response should be informative for choosing which drug might be more effective in which tumors. To evaluate our ability to choose which chemotherapy agent, docetaxel or carboplatin, would have the best response in each PDX, we used a Leave One Out Cross-Validation (LOOCV) approach to predict the quantitative response of each PDX to both carboplatin and docetaxel.

Across all the PDX models, the accuracy in predicting the best response was 69% (*Figure 6C*). However, we are particularly interested in predicting the differential response in PDXs with large differences in their carboplatin and docetaxel response. By limiting our scope to PDXs whose predicted responses to both agents differed by a log2 fold-change of more than two, the accuracy of our predictions rose to 78% (*Figure 6D*). Thus, using this approach, we can 1) identify the subset of models for which an accurate prediction can be made, and 2) predict the best therapy for these models, both with good precision.

### Preclinical Validation: Multigene Biomarker Panels Are Predictive For Chemotherapy Response In Other PDX Datasets

To test the predictive power of both our GLMs and our gene panels, we tested our predictive models on other PDX data sets (*Table 1)* with either associated RNAseq (within platform) or Affymetrix array (across platform) gene expression data. With respect to within platform validation, the carboplatin and docetaxel GLMs were tested in a PDX dataset with RNAseq and qualitative drug response from the Rosalind & Morris Goodman Cancer Research Centre (GSE142767)(Savage et al., 2020). Of the PDXs in this dataset (n=37), 30 had RECIST-like single agent response data for cisplatin and paclitaxel. Despite the differences in the PDX composition and the specific therapies used (cisplatin vs. carboplatin and paclitaxel vs. docetaxel), we found that our multigene biomarker panels were predictive in this dataset (AUC 0.60-0.67). Using the GLM for carboplatin built on our dataset, we found that AUCs for cisplatin response in this dataset were 0.67 for CR vs. all else, and 0.60 for CR/PR vs. all else (*Figure 7A*, *Figure 7B*). Similarly, when we applied the docetaxel GLM built on our dataset, the AUCs for paclitaxel response were 0.60 for CR vs. all else, and 0.67 for CR/PR vs. all else (*Figure 7A*, *Figure 7B*). Thus, our biomarker panels had predictive power for therapies that were in the same class (taxane vs. platinum-based).

**Figure 7:**
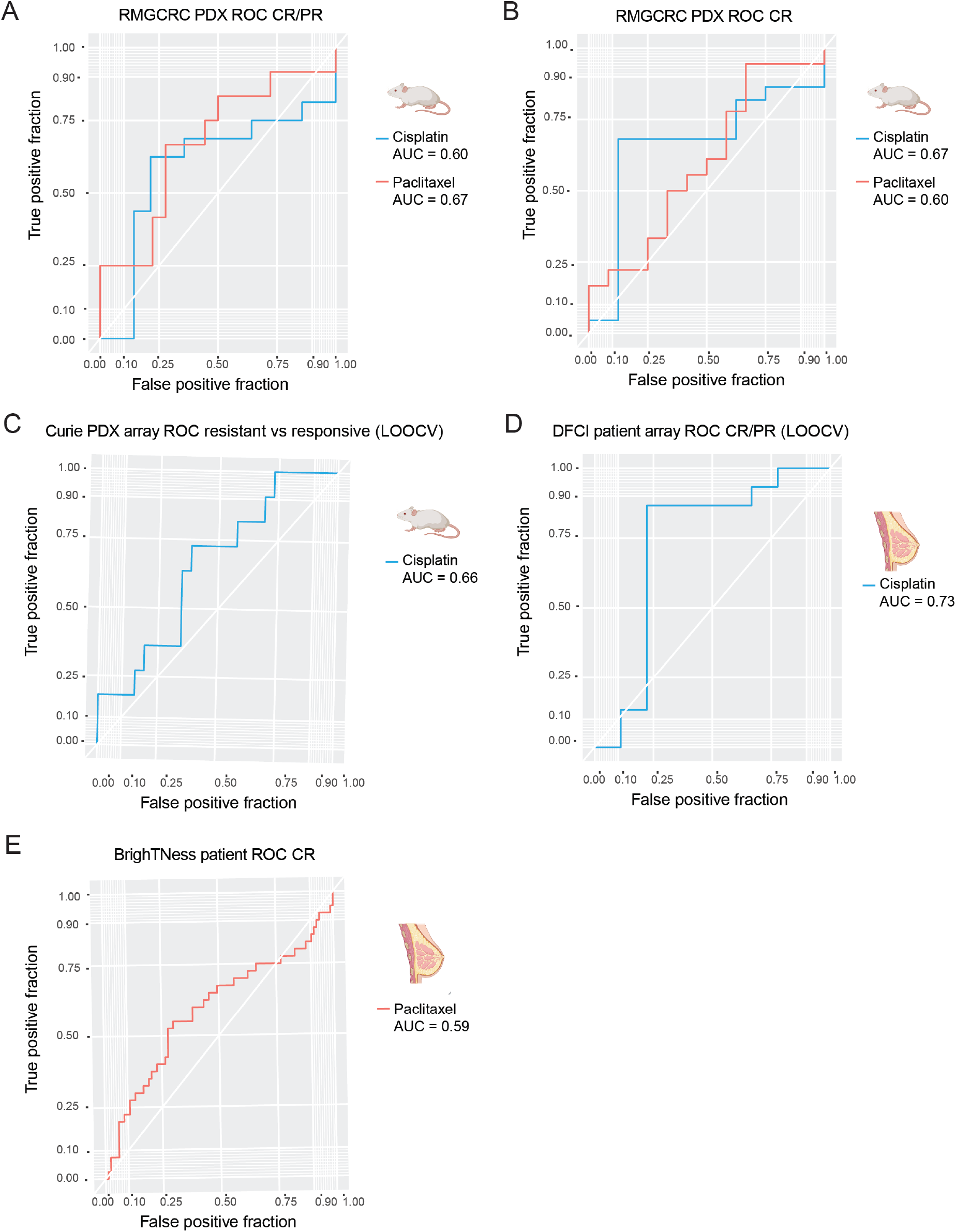
Multigene biomarkers are predictive of qualitative response in other PDX datasets and in a clinical human data set with response to cisplatin. Figure 7A: ROC of response predictions in the RMGCRC PDX cohort for complete and partial response (CR/PR) vs. all else. The ROC for predictions for cisplatin is in blue, and the ROC for predictions to paclitaxel is red. Figure 7B: ROC of response predictions in the RMGCRC PDX cohort for complete response (CR) vs. all else. The ROC for predictions for cisplatin is in blue, and the ROC for predictions to paclitaxel is red. Figure 7C: ROC of response prediction for cisplatin response in the Curie cohort. Figure 7D: ROC of response prediction for cisplatin response in a human clinical cohort (DFCI).

To test the cross-platform performance of our biomarker panel, the predictive power of our carboplatin biomarker was tested in a gene expression Affymetrix array dataset from the Curie Institute (n = 37) evaluating qualitative responses following treatment with cisplatin. Due to the platform differences in between array data, and the RNAseq data from which the GLMs were built, this dataset was used to validate the carboplatin informative genes, but not the carboplatin GLM itself. To determine the predictive power of the biomarkers, an XGBoost tree model was built with LOOCV over the 9 informative genes for carboplatin. The AUC of this model was 0.66 (*Figure* 7C). Thus, by leveraging other publicly available PDX datasets, we have shown that the multigene panels are predictive across preclinical datasets and across gene expression platforms.

### Clinical Validation: Predictive Power Of The Platinum Multigene Biomarker Panel

To investigate the predictive power of our PDX-based platinum response signature in a clinical dataset, we tested our carboplatin GLM on an array dataset with 24 TNBC patient samples (GSE18864)(Silver et al., 2010). These patients were treated with a cisplatin monotherapy and had response to therapy measured qualitatively with the RECIST 1.1 classification system. For this analysis we also used an XGBoost tree model (LOOCV), this time using the 9 informative genes for carboplatin response. The AUC for this model was 0.73 (*Figure* 7*D*), demonstrating that we have predictive power in a clinical dataset for platinum response.

To determine if the expression of stromal genes indeed contributed to the predictive power of the clinical model, we determined the variable importance rankings for this model using the varImp function from caret. The stromal genes were ranked second (*DGKQ*), third (*ADAMTS20*), fifth (*SEC14L2*), and ninth (SERPINI1) out of the nine genes for importance. This suggests that both stromal genes and epithelial genes contributed predictive power to our multigene biomarker panel.

### Clinical Validation: Predictive Power Of The Taxane Multigene Biomarker Panel

We also tested the predictive power of our taxane GLM and biomarker panel by evaluating the docetaxel GLM on a subset of TNBC patient samples from the BrighTNess clinical trial who were treated with paclitaxel. While full RECIST 1.1 calls were not available, patients were evaluated for pathologic complete response vs. not (GSE164458) (Loibl et al., 2018). Although the GLM was built using PDX models showing fewer CR relative to the number observed for carboplatin, as well as using a different taxane (docetaxel vs. paclitaxel), we still had predictive power in this clinical dataset (AUC = 0.59) (*Figure 7E*). The contribution of the stromal genes to the taxane prediction in the BrighTNess trial was also evaluated using the variable importance rankings function from caret. We found that here too the stromal genes contributed to the model and were ranked first (*PARPBP*) and fourth (*KIF24*) out of the six genes for informativeness.

## Discussion

### PDX models represent a wide range of TNBC patients and recapitulate tumor of origin responses, to docetaxel, making them a viable system to identify biomarkers of response and resistance

PDX models have emerged as useful preclinical tools. Unlike cell line models, PDXs have the advantage of modeling the patient tumor epithelium and its interaction with the stromal microenvironment, albeit mouse-derived. Herein, we demonstrate that the cohort of TNBC PDXs used in this study are highly representative of two widely used cohorts human tumors (TCGA and CPTAC) with respect to in copy number alteration patterns, mutation frequency, and gene expression profiles at both the transcriptomic and proteomic levels. Further, we demonstrate that, at least for taxanes, the responses of PDX to docetaxel are qualitatively similar to the responses of the tumors-of-origin in the patient when treated with docetaxel or paclitaxel. This high degree of concordance allowed us to treat the TNBC as a preclinical cohort to develop multigene biomarker panels predictive of differential response to two commonly used first line agents: platinums (carboplatin and cisplatin) and taxanes (docetaxel and paclitaxel).

### The Addition Of CTD To The Standard WGCNA Approach Is A Methodological Advancement That Allows For The Identification Of Multigene Biomarker Panels Associated With A Phenotype Of Interest

To identify our gene biomarker panels, we first had to address a methodological issue. The initial gene modules identified by WGCNA were large, and it was difficult to determine which specific genes were contributing to the modules being associated with response. By adding CTD to the standard WGCNA approach we identified highly connected sets of genes and identified specific genes that contributed to the predictive power of the smaller submodules. The biomarker panels identified by this approach outperformed several methods that are commonly used for feature selection including correlation analysis, a recursive feature extraction approach, and the hub genes identified with the standard WGCNA approach. Additionally, a literature search shown that many of our genes have been shown to play a role in cancer progression or resistance to therapy.

Within the carboplatin and docetaxel response prediction panels, we identified both stromal and epithelial genes whose differential expression is predictive of differential response to both classes of chemotherapy agents. Although murine stromal genes were identified as part of the original biomarker panels, their human homologs were shown to be predictive in the clinical cohort. The identification of predictive stromal genes in this analysis is also supported by studies that have previously associated breast cancer stromal cells with both progression and resistance (Conklin and Keely, 2012; Criscitiello et al., 2014; Dittmer and Leyh, 2015; Farmer et al., 2009; Plava et al., 2019).

### Multigene biomarkers proved predictive across expression platforms and chemotherapy agent classes

By testing our multigene biomarkers across datasets with different platforms and agents, we determined that 1) our multigene biomarkers are predictive in both RNAseq and microarray datasets and 2) the multigene biomarkers developed for carboplatin and docetaxel are predictive for other chemotherapy agents of the same class (platinums and taxanes, respectively). A subset of the informative genes identified in this study were also identified with text mining and with proteomics data. Interestingly, although the protein dataset was computationally separated to only include proteins that were expressed in human cells, one of the proteins that was correlated with response, DGKQ, was identified in the stromal carboplatin network. This suggest that perhaps this gene/protein is informative when it is expressed in both the murine stromal and human cancer cells.

While our study focused on identifying biomarkers of chemotherapy response, CTD can be applied to other data types and has previously been used to identify patterns of metabolic perturbations (Thistlethwaite et al., 2021). In future analyses, CTD could be applied to multiple data types to create a multi-omics signature of resistance. Our approach could also be applied to any biological problem that uses highly heterogenous, or highly dimensional data to identify specific biomarkers related to any phenotype of interest.

## Acknowledgements

This work was supported by NIH/NCI grants U54 CA224076 (to A.L.W, B.E.W, and M.T.L.), U24 CA226110, (to M.T.L.), P50 CA186784 (to C.K.O and M.T.L.), Dan L. Duncan Cancer Center (P30 Cancer Center Support Grant NCI-CA125123) (to C.K.O.), and a grant from the Common Fund of the National Institutes of Health (NIH) (5U54 DA036134) (to A.M), and by grant T32 CA203690 from the Translational Breast Cancer Research Training Program (to J.T.L.). This work was also supported by a Core Facility grant from the Cancer Prevention and Research Institute of Texas (CPRIT Core Facilities Support Grant RP170691), a grant from the V Foundation, and a generous gift from the Korell family for the study of triple-negative breast cancer.The BCM Mass Spectrometry Proteomics Core is supported by the Dan L. Duncan Comprehensive Cancer Center NIH award (P30 CA125123), and a CPRIT Core Facility Award (RP170005),

## Methods

### Patient Derived Xenograft Model Selection and Chemotherapy Response

Information related to publicly available PDX models from Baylor College of Medicine (BCM) and the Huntsman Cancer Institute (HCI) can be found on the BCM PDX portal (https://pdxportal.research.bcm.edu/). Single agent docetaxel, carboplatin and untreated/vehicle control data were derived from three separate preclinical trials (to be described in full elsewhere). In total, 50 TNBC PDX models were treated across these three preclinical trials. To define a homogenous set of tumors that did not over-represent any particular group of patient samples, we eliminated PDX models that did not match an immunosuppressed basal profile or were derived from the same patient as other models in this collection. Ultimately, 45 PDX were used for the generation of predictive models this study (Supplemental Table 3).

Each of the three preclinical trials used the same treatment strategy. Briefly, PDX-bearing mice were treated with four weekly cycles of docetaxel (20mg/kg, IP) (low human equivalent dose), carboplatin (50mg/kg, IP) (human equivalent dose AUC6), or left untreated for four weeks. A subset of models also received docetaxel at 30mg/kg (high human-equivalent dose). In addition, a subset of PDX models for whom we had clinical responses of the tumor-of-origin were also treated with doxorubicin (3mg/kg)/cyclophophamide (100mg/kg) (AC), but human-equivalent doses could not be achieved under the conditions used.

Tumor volume was assessed twice weekly using calipers. Treatment responses were evaluated quantitatively by the log2 fold-change in tumor volume relative to their volume prior to treatment (~ 200mm3). A qualitative “best clinical response” was assigned using a modified RECIST 1.1 criteria allowing for more direct comparison with tumor response in patients. Complete response (CR) was defined as non-palpable. Partial response (PR) was defined as those PDX showing >30% reduction in tumor volume, but not reaching the non-palpable state. PDX lines showing stable disease (SD) showed a <30% decrease, and no more than a 20% increase over starting volume. PDX lines showing progressive disease (PD) increased > 20% over starting volume vs. vehicle control at the end of the study. Quantitative response is summarized in Supplemental Table 3.

### Genomic Sequencing

DNA sequencing was performed by core facilities at BCM and Cornell University. Exome Sequencing Libraries were prepared using the Agilent SureSelect XT v6.0 Human kit. Sequencing was performed on an Illumina HiSeq 4000 machine (PE, 2×100 cycles, ~133X estimated coverage per sample) at Cornell. At BCM, sequencing was performed on Illumina NovaSeq machine (PE, 2×100, ~200M read pairs per sample). Separation of human and mouse reads, alignment to the reference genome, variant calling, and annotation were performed using the PDXNet Tumor-Only Variant Calling pipeline developed by the Jackson Laboratory and hosted on the Cancer Genomics Cloud.

### Baseline RNAseq Transcriptomics

For RNA extraction from untreated PDX tissue, snap frozen tissue from an early transplant generation (e.g. TG1-TG5) xenograft-derived tumors were harvested and stored at −80 °C prior to use. For library preparation, the NuGEN Ovation RNAseq v2 Kit (protocol p/n 7102, reagent kit p/n 7102-08) along with ThermoFisher’s ERCC RNA Spike-In Control Mixes Protocol (publication number 4455352, rev. D) and either the Illumina TruSeq DNA-Seq (protocol p/n 15005180 Rev. C, June 2011, reagent kit p/n FC-121-2001) or the Rubicon ThruPlex DNA-Seq (protocol: QAM-108-002, kit p/n R400428) protocols were used.

FastQ file generation was executed using bcl2fast software or Illumina's cloud-based informatics platform, BaseSpace Sequencing Hub.

### Baseline Mass Spectrometry-based Proteomics

The mass spectrometry proteomic iBAQ values were obtained for the same set of PDX models. To identify proteins that are specifically expressed in the human cancer cells, gpGrouper was used to deconvolute murine vs human proteins. The iBAQ values were then used to identify differentially expressed proteins.

### Bioinformatics Analyses And Comparisons With TCGA And CPTAC Samples

#### DNA Sequencing Data Processing

All raw FASTQ files were subjected to QC verification by FASTQC (https://www.bioinformatics.babraham.ac.uk/projects/fastqc/) and were trimmed for adapter sequences with TrimGalore (https://www.bioinformatics.babraham.ac.uk/projects/trim_galore/). Whole Exome FASTQ files were processed using the Tumor-Only Variant Calling pipeline (Evrard et al., 2019) developed by the Jackson Laboratory for PDXNet hosted on the Cancer Genomics Cloud. This pipeline uses Xenome(Conway et al., 2012) to separate human epithelial reads and mouse stromal reads, then aligns human reads against the GRCh38 human genome using bwa (Li, 2013). It then uses GATK MuTect2 (DePristo et al., 2011; Marx, 2013; McKenna et al., 2010) to call variants and SnpEff/SnpSift (Cingolani et al., 2012a, 2012b) to annotate variants, generating per-sample VCF and tabular outputs for downstream analyses. Further annotation with gnomAD (Karczewski et al., 2020) and CLINVAR (Landrum et al., 2017) was performed which added information by matching locus fields (Chromosome, Position, Reference and Alternate alleles) to standard annotation VCF files. Multi-allelic sites were decomposed to ensure Alternate alleles matched exactly with annotation sources.

#### Copy Number Analysis

A set of 6 normal breast tissue samples were subjected to Whole Exome Sequencing both at Cornell and Baylor to serve as platform-matched normal samples. These platforms matched normal samples were run through the Variant Calling pipeline to obtain platform-matched normal BAMs. CopywriteR(Kuilman et al., 2015) was used on each tumor BAM file obtained from the Variant Calling pipeline, and a random platform-matched normal BAM was used as control. CopywriteR uses depth information from off-target reads to calculate segment-level copy number data. This segment level data was then processed using GISTIC2.0(Mermel et al., 2011) to obtain raw and threshold gene-level and focal-level copy number information.

#### Comparison with TCGA Samples

Mutation, Copy Number and Clinical Biomarker data for TCGA-BRCA samples was obtained using the TCGABiolinks package. TCGA-BRCA samples and PDX models were grouped based on their ER/PR and HER2 IHC status, and each TCGA group was compared with its equivalent group of PDX models.

Mutation frequency was computer per gene for all TCGA and PDX groups and compared among TNBC, ER+ and HER2+ groups. Similarly, Copy Number segment data was plotted per group to visualize common patterns in each group. To examine similarities between TNBC TCGA samples and TNBC PDX models, the segment level data for all TNBC samples across the groups were analyzed with GISTIC2.0 and the focal level copy number changes were used to generate a distance matrix and dendrogram using the R packages ggplot2(Wickham, 2016),dendextend(Galili, 2015) and circlize (Gu et al., 2014). A similar analysis was also done for ER+ and HER2+ groups.

#### Gene Expression Quantification

RNAseq FASTQ files were processed using Xenome to separate murine stromal reads and human epithelial reads, which were both then quantified with rsem-calculate-expression. A reference index was created for RSEM (v1.3.0) (Li and Dewey, 2011) using rsem-prepare-reference on the hg38 and mm10 genome assembly (FASTA) and transcript feature (GTF) files. Expected counts from RSEM were collated to create a genes-by-samples matrix. This matrix was Upper-Quantile normalized per-sample to account for differences in sequencing depth.

#### Network Building

To identify genes that could be predictive of response to chemotherapy, we used a gene network approach. For both the human and mouse reads, a network was built over all of the PDXs and the responsive PDXs for both carboplatin and docetaxel independently. The gene expression data was obtained from deep RNAseq (~200M reads/sample) of the PDXs. Preprocessing included removing genes that were expressed in less than 20% of the samples or had a low expression across all of the samples. The five thousand most variable genes were then selected for the human and mouse reads. These genes were used to build the species-specific networks. The following graphs were built: an epithelial graph with all the PDXs, an epithelial graph with the samples that are responsive to carboplatin, an epithelial graph with the samples that are responsive to docetaxel, a stromal graph from all the samples, a stromal graph with the samples that are responsive to carboplatin, and a stromal graph with the samples that are responsive to docetaxel. The graphs were then pruned to remove weak edges (defined as having an edge weight of <0.2). The graphs built from all the samples were then pruned to remove the edges that were also found in the responsive graphs. This was done for both carboplatin and docetaxel in both the epithelial and stromal compartments to identify signatures linked to resistance and potentially resistance mechanisms.

#### Module Discovery and the Application of CTD

Coarse modules were defined in the pruned graphs by applying WGCNA’s cutreeDynamic function. These modules were large, so we used CTD(Thistlethwaite et al., 2021) to identify highly connected genes within the modules. This allowed us to identify smaller and more biologically tractable sets of genes. We then determined which of these submodules and which specific genes within these submodules were informative for resistance.

#### Text Mining

To determine if our method selected genes associated with for chemotherapy resistance genes in comparison to the standard WGCNA approach, we utilized a text mining tool, iTextMine(Ren et al., 2018)

#### Predicting response to chemotherapy

The genes identified as informative were used to predict the response to carboplatin and docetaxel and the combination treatment. To model bulk patient data better, a pseudo-bulk profile was first made for each of the PDXs in our collection by summing the human epithelial reads and the murine stromal reads. Ten balanced 30% test and 70% training sets were defined using caret’s (Kuhn, 2008)createDataPartition function for each treatment. For each of these training/test permutations, the training set was used to create a quantitative GLM, whose performance was then tested on the test set. The mean absolute error and root mean square error were calculated for each training/test permutation and the observed vs. predicted log2 (fold change) was plotted. An LOOCV approach was used to determine the predictive power of our model in determining resistance vs. response as defined by RECIST criteria. The quantitative response for each sample was predicted, then the AUC was calculated to determine based on the qualitative response.

#### Comparison of our WGCNA/CTD approach to other feature selection methods

To compare the MAE and RMSE of the WGCNA/CTD approach to other commonly used feature selection methods, we also choose potentially informative genes with three other commonly used feature selection methods (correlation, recursive feature extraction, hub genes from informative WGCNA modules). Correlation - for each treatment the top 10 most correlated epithelial and stromal genes were determined. Recursive feature extraction – for each treatment the top 10 most informative epithelial and stromal genes were determined. Hub genes – to identify the hub genes for each treatment, WGCNA modules whose eigengene was correlation with response > 0.35 were identified. Then genes with a >0.75 correlation with the module eigengene and a >0.2 correlation with response were identified as the hub genes. For carboplatin there were 14 epithelial WGCNA hub genes and no stromal WGCNA hub genes. For docetaxel there were 26 epithelial WGCNA genes and 1 stromal WGCNA gene.

#### Rosalind & Morris Goodman Cancer Research Centre RNAseq PDX dataset prediction

An RNAseq dataset and qualitative response were obtained from GEO (GSE142767) (n = 37). The majority of these PDX models were Basal-like and triple negative, however there were some PDX models that were classified as HER2(7) or positive for ER (5) or the HER2 enrichment (3). A GLM was built over all 45 PDXs in our collection over the informative genes for carboplatin, and this GLM was then applied to the validation RNAseq dataset. A ROC curve was then calculated to determine the predictive power of the GLM in predicting if the PDX has CR vs. all else or CR and PR vs. all else.

#### Curie dataset prediction

The Curie dataset was obtained from the Curie institute. This dataset consists of the Affymetrix array expression of 37 TNBC PDXs and the human equivalent response to cisplatin. Due to the inconsistencies between the array and RNAseq platforms, an XGBoost tree model (LOOCV) was built over the carboplatin multigene biomarker with the xgbTree option in caret for this dataset to determine the AUC and predictive nature of these genes.

#### Dana-Farber Cancer Institute TNBC patient array dataset prediction

The Dana-Farber (GSE18864) dataset was obtained from GEO (Silver et al., 2010). This dataset included array data from 24 patient biopsies as well as qualitative response of these patients to neoadjuvant cisplatin. These patients were part of a cohort of 28 women with TNBC that were treated with neoadjuvant cisplatin every 21 days. An XGBoost tree model (LOOCV) was built over the carboplatin multigene biomarker with the xgbTree option in caret for this dataset to determine the AUC and predictive nature of these genes.

#### BrighTNess TNBC patient RNAseq dataset prediction

RNAseq and patient response to paclitaxel was obtained for the BrighTNess (GSE164458) dataset from GEO (Loibl et al., 2018). Data from arm C, single agent paclitaxel in 123 patients were used for this study. A GLM was built for the PDXs in our collection over the informative genes for docetaxel, and this GLM was then used to predict the response (pCR or not) to paclitaxel.

## Notes

**Disclosures** M.T.L is a Founder of, and an uncompensated Manager in StemMed Holdings L.L.C., an uncompensated Limited Partner in StemMed Ltd., and is a Founder of and equity stake holder in Tvardi Therapeutics. L.D. is a compensated employee of StemMed Ltd. Selected BCM PDX models described herein are exclusively licensed to StemMed Ltd. resulting in tangible property royalties to M.T.L. and L.D. University of Utah may license the HCI PDX models described herein to for-profit companies, which may result in tangible property royalties to A.L.W. and B.E.W. Washington University has licensed selected PDX to Envigo which results in tangible property royalties to S.L. He also received research funding from Pfizer, Takeda Oncology, Zenopharm, independent of this project. S.L has received license fees from Envigo. He received research funding from Pfizer, Takeda Oncology, Zenopharm, outside of this project. O.S is a compensated employee of, and equity stake holder in, Bluebird Bio.

### Competing Interest Statement

M.T.L is a Founder of, and an uncompensated Manager in StemMed Holdings L.L.C., an uncompensated Limited Partner in StemMed Ltd., and is a Founder of and equity stake holder in Tvardi Therapeutics. L.D. is a compensated employee of StemMed Ltd. Selected BCM PDX models described herein are exclusively licensed to StemMed Ltd. resulting in tangible property royalties to M.T.L. and L.D. University of Utah may license the HCI PDX models described herein to for-profit companies, which may result in tangible property royalties to A.L.W. and B.E.W. Washington University has licensed selected PDX to Envigo which results in tangible property royalties to S.L. He also received research funding from Pfizer, Takeda Oncology, Zenopharm, independent of this project. S.L has received license fees from Envigo. He received research funding from Pfizer, Takeda Oncology, Zenopharm, outside of this project. O.S is a compensated employee of, and equity stake holder in, Bluebird Bio.

## References

Bai, X., Gao, C., Zhang, L., and Yang, S. (2019). Integrin α7 high expression correlates with deteriorative tumor features and worse overall survival, and its knockdown inhibits cell proliferation and invasion but increases apoptosis in breast cancer. J Clin Lab Anal 33, e22979.

Bandala, C., Cortés-Algara, A.L., Mejía-Barradas, C.M., Ilizaliturri-Flores, I., Dominguez-Rubio, R., Bazán-Méndez, C.I., Floriano-Sánchez, E., Luna-Arias, J.P., Anaya-Ruiz, M., and Lara-Padilla, E. (2015). Botulinum neurotoxin type A inhibits synaptic vesicle 2 expression in breast cancer cell lines. Int J Clin Exp Patho 8, 8411–8418.

Bhandari, A., Xia, E., Zhou, Y., Guan, Y., Xiang, J., Kong, L., Wang, Y., Yang, F., Wang, O., and Zhang, X. (2018). ITGA7 functions as a tumor suppressor and regulates migration and invasion in breast cancer. Cancer Management Res 10, 969–976.

Bozza, W.P., Zhang, Y., Hallett, K., Rosado, L.A.R., and Zhang, B. (2015). RhoGDI deficiency induces constitutive activation of Rho GTPases and COX-2 pathways in association with breast cancer progression. Oncotarget 6, 32723–32736.

Chen, B., Lai, J., Dai, D., Chen, R., Liao, N., and Tang, H. (2020). PARPBP is a prognostic marker and confers chemotherapeutic resistance to breast cancer.

Cingolani, P., Platts, A., Wang, L.L., Coon, M., Nguyen, T., Wang, L., Land, S.J., Lu, X., and Ruden, D.M. (2012a). A program for annotating and predicting the effects of single nucleotide polymorphisms, SnpEff. Fly 6, 80–92.

Cingolani, P., Patel, V.M., Coon, M., Nguyen, T., Land, S.J., Ruden, D.M., and Lu, X. (2012b). Using Drosophila melanogaster as a Model for Genotoxic Chemical Mutational Studies with a New Program, SnpSift. Frontiers Genetics 3, 35.

Conklin, M.W., and Keely, P.J. (2012). Why the stroma matters in breast cancer: insights into breast cancer patient outcomes through the examination of stromal biomarkers. Cell Adhes Migr 6, 249–260.

Conway, T., Wazny, J., Bromage, A., Tymms, M., Sooraj, D., Williams, E.D., and Beresford-Smith, B. (2012). Xenome—a tool for classifying reads from xenograft samples. Bioinformatics 28, i172–i178.

Criscitiello, C., Esposito, A., and Curigliano, G. (2014). Tumor–stroma crosstalk: targeting stroma in breast cancer. Curr Opin Oncol 26, 551–555.

Dany, M., and Ogretmen, B. (2015). Ceramide induced mitophagy and tumor suppression. Biochimica Et Biophysica Acta Bba - Mol Cell Res 1853, 2834–2845.

Darb-Esfahani, S., Kronenwett, R., Minckwitz, G. von, Denkert, C., Gehrmann, M., Rody, A., Budczies, J., Brase, J.C., Mehta, M.K., Bojar, H., et al. (2012). Thymosin beta 15A (TMSB15A) is a predictor of chemotherapy response in triple-negative breast cancer. Brit J Cancer 107, 1892–1900.

DePristo, M.A., Banks, E., Poplin, R., Garimella, K.V., Maguire, J.R., Hartl, C., Philippakis, A.A., Angel, G. del, Rivas, M.A., Hanna, M., et al. (2011). A framework for variation discovery and genotyping using next-generation DNA sequencing data. Nat Genet 43, 491–498.

Di, Y., Chen, D., Yu, W., and Yan, L. (2019). Bladder cancer stage-associated hub genes revealed by WGCNA co-expression network analysis. Hereditas 156, 7.

Dittmer, J., and Leyh, B. (2015). The impact of tumor stroma on drug response in breast cancer. Semin Cancer Biol 31, 3–15.

Dobrolecki, L.E., Airhart, S.D., Alferez, D.G., Aparicio, S., Behbod, F., Bentires-Alj, M., Brisken, C., Bult, C.J., Cai, S., Clarke, R.B., et al. (2016). Patient-derived xenograft (PDX) models in basic and translational breast cancer research. Cancer Metast Rev 35, 547–573.

Du, B., Yuan, L., Sun, L., and Zhang, Z. (2020). WGCNA screening of prognostic markers in medulloblastoma. National Medical J China 100, 460–464.

Echeverria, G.V., Powell, E., Seth, S., Ge, Z., Carugo, A., Bristow, C., Peoples, M., Robinson, F., Qiu, H., Shao, J., et al. (2018). High-resolution clonal mapping of multi-organ metastasis in triple negative breast cancer. Nat Commun 9, 5079.

Evrard, Y.A., Srivastava, A., Randjelovic, J., Arunachalam, S., Bult, C.J., Chen, H., Chen, L., Davies, M., Davies, S., Davis-Dusenbery, B., et al. (2019). Systematic Establishment of Robustness and Standards in Patient-Derived Xenograft Experiments and Analysis. Biorxiv 790246.

Farmer, P., Bonnefoi, H., Anderle, P., Cameron, D., Wirapati, P., Wirapati, P., Becette, V., André, S., Piccart, M., Campone, M., et al. (2009). A stroma-related gene signature predicts resistance to neoadjuvant chemotherapy in breast cancer. Nat Med 15, 68–74.

Galili, T. (2015). dendextend: an R package for visualizing, adjusting and comparing trees of hierarchical clustering. Bioinformatics 31, 3718–3720.

Garrido-Castro, A.C., Lin, N.U., and Polyak, K. (2019). Insights into Molecular Classifications of Triple-Negative Breast Cancer: Improving Patient Selection for Treatment. Cancer Discov 9, 176–198.

Giulietti, M., Occhipinti, G., Principato, G., and Piva, F. (2017). Identification of candidate miRNA biomarkers for pancreatic ductal adenocarcinoma by weighted gene co-expression network analysis. Cell Oncol 40, 181–192.

Gu, Z., Gu, L., Eils, R., Schlesner, M., and Brors, B. (2014). circlize implements and enhances circular visualization in R. Bioinformatics 30, 2811–2812.

Gu, Z., Eils, R., and Schlesner, M. (2016). Complex heatmaps reveal patterns and correlations in multidimensional genomic data. Bioinformatics 32, 2847–2849.

Guo, X., Li, J., Zhang, H., Liu, H., Liu, Z., and Wei, X. (2018). Relationship Between ADAMTS8, ADAMTS18, and ADAMTS20 (A Disintegrin and Metalloproteinase with Thrombospondin Motifs) Expressions and Tumor Molecular Classification, Clinical Pathological Parameters, and Prognosis in Breast Invasive Ductal Carcinoma. Medical Sci Monit Int Medical J Exp Clin Res 24, 3726–3735.

Huang, Y., Liu, H., Zuo, L., and Tao, A. (2020). Key genes and co-expression modules involved in asthma pathogenesis. Peerj 8, e8456.

Izumchenko, E., Paz, K., Ciznadija, D., Sloma, I., Katz, A., Vasquez-Dunddel, D., Ben-Zvi, I., Stebbing, J., McGuire, W., Harris, W., et al. (2017). Patient-derived xenografts effectively capture responses to oncology therapy in a heterogeneous cohort of patients with solid tumors. Ann Oncol 28, 2595–2605.

Jia, B., Zhao, X., Wang, Y., Wang, J., Wang, Y., and Yang, Y. (2019). Prognostic roles of MAGE family members in breast cancer based on KM-Plotter Data. Oncol Lett 18, 3501–3516.

Jia, R., Zhao, H., and Jia, M. (2020). Identification of co-expression modules and potential biomarkers of breast cancer by WGCNA. Gene 750, 144757.

Karczewski, K.J., Francioli, L.C., Tiao, G., Cummings, B.B., Alföldi, J., Wang, Q., Collins, R.L., Laricchia, K.M., Ganna, A., Birnbaum, D.P., et al. (2020). The mutational constraint spectrum quantified from variation in 141,456 humans. Nature 581, 434–443.

Kim, S., Lee, K., Choi, J.-H., Ringstad, N., and Dynlacht, B.D. (2015). Nek2 activation of Kif24 ensures cilium disassembly during the cell cycle. Nat Commun 6, 8087.

Koboldt, D.C., Fulton, R.S., McLellan, M.D., Schmidt, H., Kalicki-Veizer, J., McMichael, J.F., Fulton, L.L., Dooling, D.J., Ding, L., Mardis, E.R., et al. (2012). Comprehensive molecular portraits of human breast tumours. Nature 490, 61–70.

Kuhn, M. (2008). Building Predictive Models in R Using the caret Package. Wiley Interdiscip Rev Comput Statistics 1, 128–129.

Kuilman, T., Velds, A., Kemper, K., Ranzani, M., Bombardelli, L., Hoogstraat, M., Nevedomskaya, E., Xu, G., Ruiter, J. de, Lolkema, M.P., et al. (2015). CopywriteR: DNA copy number detection from off-target sequence data. Genome Biol 16, 49.

Lagadec, C., Vlashi, E., Frohnen, P., Alhiyari, Y., Chan, M., and Pajonk, F. (2014). The RNA-binding protein Musashi-1 regulates proteasome subunit expression in breast cancer- and glioma-initiating cells. Stem Cells Dayt Ohio 32, 135–144.

Landrum, M.J., Lee, J.M., Benson, M., Brown, G.R., Chao, C., Chitipiralla, S., Gu, B., Hart, J., Hoffman, D., Jang, W., et al. (2017). ClinVar: improving access to variant interpretations and supporting evidence. Nucleic Acids Res 46, gkx1153-.

Lang, Y., Kong, X., He, C., Wang, F., Liu, B., Zhang, S., Ning, J., Zhu, K., and Xu, S. (2017). Musashi1 Promotes Non-Small Cell Lung Carcinoma Malignancy and Chemoresistance via Activating the Akt Signaling Pathway. Cell Physiology Biochem Int J Exp Cell Physiology Biochem Pharmacol 44, 455–466.

Li, H. (2013). Aligning sequence reads, clone sequences and assembly contigs with BWA-MEM. Arxiv.

Li, B., and Dewey, C.N. (2011). RSEM: accurate transcript quantification from RNA-Seq data with or without a reference genome. Bmc Bioinformatics 12, 323.

Liao, Y., Wang, Y., Cheng, M., Huang, C., and Fan, X. (2020). Weighted Gene Coexpression Network Analysis of Features That Control Cancer Stem Cells Reveals Prognostic Biomarkers in Lung Adenocarcinoma. Frontiers Genetics 11, 311.

Liu, W., Li, L., Ye, H., and Tu, W. (2017). [Weighted gene co-expression network analysis in biomedicine research]. Sheng Wu Gong Cheng Xue Bao Chin J Biotechnology 33, 1791–1801.

Loibl, S., O’Shaughnessy, J., Untch, M., Sikov, W.M., Rugo, H.S., McKee, M.D., Huober, J., Golshan, M., Minckwitz, G. von, Maag, D., et al. (2018). Addition of the PARP inhibitor veliparib plus carboplatin or carboplatin alone to standard neoadjuvant chemotherapy in triple-negative breast cancer (BrighTNess): a randomised, phase 3 trial. Lancet Oncol 19, 497–509.

Marx, V. (2013). Genomics in the clouds. Nat Methods 10, 941–945.

McKenna, A., Hanna, M., Banks, E., Sivachenko, A., Cibulskis, K., Kernytsky, A., Garimella, K., Altshuler, D., Gabriel, S., Daly, M., et al. (2010). The Genome Analysis Toolkit: A MapReduce framework for analyzing next-generation DNA sequencing data. Genome Res 20, 1297–1303.

Mehanna, J., Haddad, F.G., Eid, R., Lambertini, M., and Kourie, H.R. (2019). Triple-negative breast cancer: current perspective on the evolving therapeutic landscape. Int J Women’s Heal Volume 11, 431–437.

Mermel, C.H., Schumacher, S.E., Hill, B., Meyerson, M.L., Beroukhim, R., and Getz, G. (2011). GISTIC2.0 facilitates sensitive and confident localization of the targets of focal somatic copy-number alteration in human cancers. Genome Biol 12, R41.

Minckwitz, G. von, Schneeweiss, A., Loibl, S., Salat, C., Denkert, C., Rezai, M., Blohmer, J.U., Jackisch, C., Paepke, S., Gerber, B., et al. (2014). Neoadjuvant carboplatin in patients with triple-negative and HER2-positive early breast cancer (GeparSixto; GBG 66): a randomised phase 2 trial. Lancet Oncol 15, 747–756.

Ming, X.-Y., Fu, L., Zhang, L.-Y., Qin, Y.-R., Cao, T.-T., Chan, K.W., Ma, S., Xie, D., and Guan, X.-Y. (2016). Integrin α7 is a functional cancer stem cell surface marker in oesophageal squamous cell carcinoma. Nat Commun 7, 13568.

Plava, J., Cihova, M., Burikova, M., Matuskova, M., Kucerova, L., and Miklikova, S. (2019). Recent advances in understanding tumor stroma-mediated chemoresistance in breast cancer. Mol Cancer 18, 67.

Powell, R.T., Redwood, A., Liu, X., Guo, L., Cai, S., Zhou, X., Tu, Y., Zhang, X., Qi, Y., Jiang, Y., et al. (2020). Pharmacologic profiling of patient-derived xenograft models of primary treatment-naïve triple-negative breast cancer. Sci Rep-Uk 10, 17899.

Qiu, J., Du, Z., Wang, Y., Zhou, Y., Zhang, Y., Xie, Y., and Lv, Q. (2019). Weighted gene co-expression network analysis reveals modules and hub genes associated with the development of breast cancer. Medicine 98, e14345.

Ren, J., Li, G., Ross, K., Arighi, C., McGarvey, P., Rao, S., Cowart, J., Madhavan, S., Vijay-Shanker, K., and Wu, C.H. (2018). iTextMine: integrated text-mining system for large-scale knowledge extraction from the literature. Database 2018.

Saltzman, A.B., Leng, M., Bhatt, B., Singh, P., Chan, D.W., Dobrolecki, L., Chandrasekaran, H., Choi, J.M., Jain, A., Jung, S.Y., et al. (2018). gpGrouper: A Peptide Grouping Algorithm for Gene-Centric Inference and Quantitation of Bottom-Up Proteomics Data*. Mol Cell Proteomics 17, 2270–2283.

Savage, P., Pacis, A., Kuasne, H., Liu, L., Lai, D., Wan, A., Dankner, M., Martinez, C., Muñoz-Ramos, V., Pilon, V., et al. (2020). Chemogenomic profiling of breast cancer patient-derived xenografts reveals targetable vulnerabilities for difficult-to-treat tumors. Commun Biology 3, 310.

Sharma, P., López-Tarruella, S., García-Saenz, J.A., Ward, C., Connor, C.S., Gómez, H.L., Prat, A., Moreno, F., Jerez-Gilarranz, Y., Barnadas, A., et al. (2017). Efficacy of Neoadjuvant Carboplatin plus Docetaxel in Triple-Negative Breast Cancer: Combined Analysis of Two Cohorts. Clin Cancer Res 23, 649–657.

Sikov, W.M., Berry, D.A., Perou, C.M., Singh, B., Cirrincione, C.T., Tolaney, S.M., Kuzma, C.S., Pluard, T.J., Somlo, G., Port, E.R., et al. (2014). Impact of the Addition of Carboplatin and/or Bevacizumab to Neoadjuvant Once-per-Week Paclitaxel Followed by Dose-Dense Doxorubicin and Cyclophosphamide on Pathologic Complete Response Rates in Stage II to III Triple-Negative Breast Cancer: CALGB 40603 (Alliance). J Clin Oncol 33, 13–21.

Silver, D.P., Richardson, A.L., Eklund, A.C., Wang, Z.C., Szallasi, Z., Li, Q., Juul, N., Leong, C.-O., Calogrias, D., Buraimoh, A., et al. (2010). Efficacy of Neoadjuvant Cisplatin in Triple-Negative Breast Cancer. J Clin Oncol 28, 1145–1153.

Spring, L.M., Fell, G., Arfe, A., Sharma, C., Greenup, R., Reynolds, K.L., Smith, B.L., Alexander, B., Moy, B., Isakoff, S.J., et al. (2020). Pathologic Complete Response after Neoadjuvant Chemotherapy and Impact on Breast Cancer Recurrence and Survival: A Comprehensive Meta-analysis. Clin Cancer Res 26, 2838–2848.

Staudigl, C., Concin, N., Grimm, C., Pfeiler, G., Nehoda, R., Singer, C.F., and Polterauer, S. (2015). Prognostic relevance of pretherapeutic gamma-glutamyltransferase in patients with primary metastatic breast cancer. Plos One 10, e0125317.

Tadist, K., Najah, S., Nikolov, N.S., Mrabti, F., and Zahi, A. (2019). Feature selection methods and genomic big data: a systematic review. J Big Data 6, 79.

Tang, J., Kong, D., Cui, Q., Wang, K., Zhang, D., Gong, Y., and Wu, G. (2018). Prognostic Genes of Breast Cancer Identified by Gene Co-expression Network Analysis. Frontiers Oncol 8, 374.

Thistlethwaite, L.R., Petrosyan, V., Li, X., Miller, M.J., Elsea, S.H., and Milosavljevic, A. (2021). CTD: An information-theoretic algorithm to interpret sets of metabolomic and transcriptomic perturbations in the context of graphical models. Plos Comput Biol 17, e1008550.

Valiente, M., Obenauf, A.C., Jin, X., Chen, Q., Zhang, X.H.-F., Lee, D.J., Chaft, J.E., Kris, M.G., Huse, J.T., Brogi, E., et al. (2014). Serpins Promote Cancer Cell Survival and Vascular Co-Option in Brain Metastasis. Cell 156, 1002–1016.

Wahba, H.A., and El-Hadaad, H.A. (2015). Current approaches in treatment of triple-negative breast cancer. Cancer Biology Medicine 12, 106–116.

Whittle, J.R., Lewis, M.T., Lindeman, G.J., and Visvader, J.E. (2015). Patient-derived xenograft models of breast cancer and their predictive power. Breast Cancer Res Bcr 17, 523.

Wickham, H. (2016). ggplot2, Elegant Graphics for Data Analysis. R 147–168.

Woo, X.Y., Giordano, J., Srivastava, A., Zhao, Z.-M., Lloyd, M.W., Bruijn, R. de, Suh, Y.-S., Patidar, R., Chen, L., Scherer, S., et al. (2019). Conservation of copy number profiles during engraftment and passaging of patient-derived cancer xenografts. Biorxiv 861393.

Zhang, B., Zhang, Y., Dagher, M.-C., and Shacter, E. (2005). Rho GDP Dissociation Inhibitor Protects Cancer Cells against Drug-Induced Apoptosis. Cancer Res 65, 6054–6062.

Zhang, L., Kang, W., Lu, X., Ma, S., Dong, L., and Zou, B. (2018). Weighted gene co-expression network analysis and connectivity map identifies lovastatin as a treatment option of gastric cancer by inhibiting HDAC2. Gene 681, 15–25.

Zhang, X., Claerhout, S., Prat, A., Dobrolecki, L.E., Petrovic, I., Lai, Q., Landis, M.D., Wiechmann, L., Schiff, R., Giuliano, M., et al. (2013a). A renewable tissue resource of phenotypically stable, biologically and ethnically diverse, patient-derived human breast cancer xenograft models. Cancer Res 73, 4885–4897.

Zhang, X., Claerhout, S., Prat, A., Dobrolecki, L.E., Petrovic, I., Lai, Q., Landis, M.D., Wiechmann, L., Schiff, R., Giuliano, M., et al. (2013b). A Renewable Tissue Resource of Phenotypically Stable, Biologically and Ethnically Diverse, Patient-Derived Human Breast Cancer Xenograft Models. Cancer Res 73, 4885–4897.

Zhao, W., Langfelder, P., Fuller, T., Dong, J., Li, A., and Hovarth, S. (2010). Weighted gene coexpression network analysis: state of the art. J Biopharm Stat 20, 281–300.

