## Supplemental Figures 1-3 and Supplemental Tables for "A Network Approach to Identify Biomarkers of Differential Chemotherapy Response Using Patient-Derived Xenografts of Triple-Negative Breast Cancer"

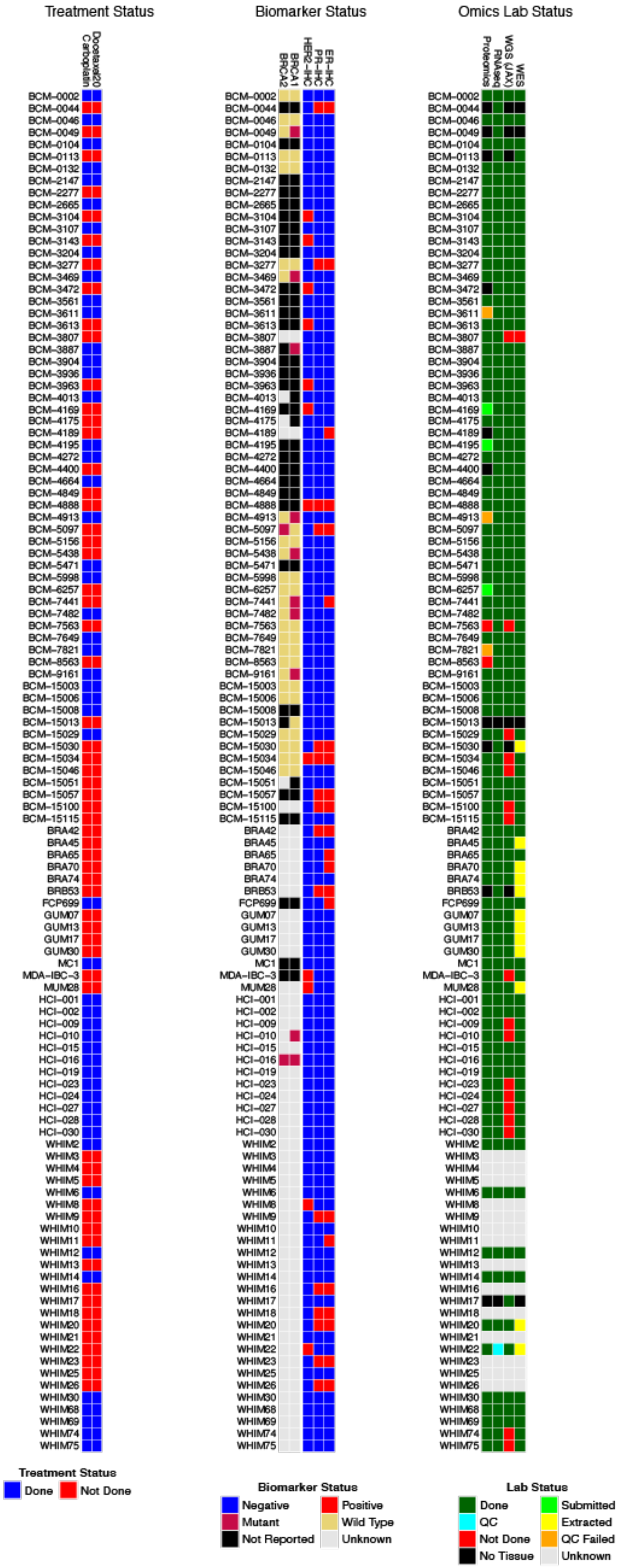

*Supplemental Figure 1: Summary of all available PDX models. Furthest to the left is the treatment status for Carboplatin and Docetaxel. The middle set of bars is the biomarker status. The rightmost set of bars is the "omics" profiling status.*

A

B

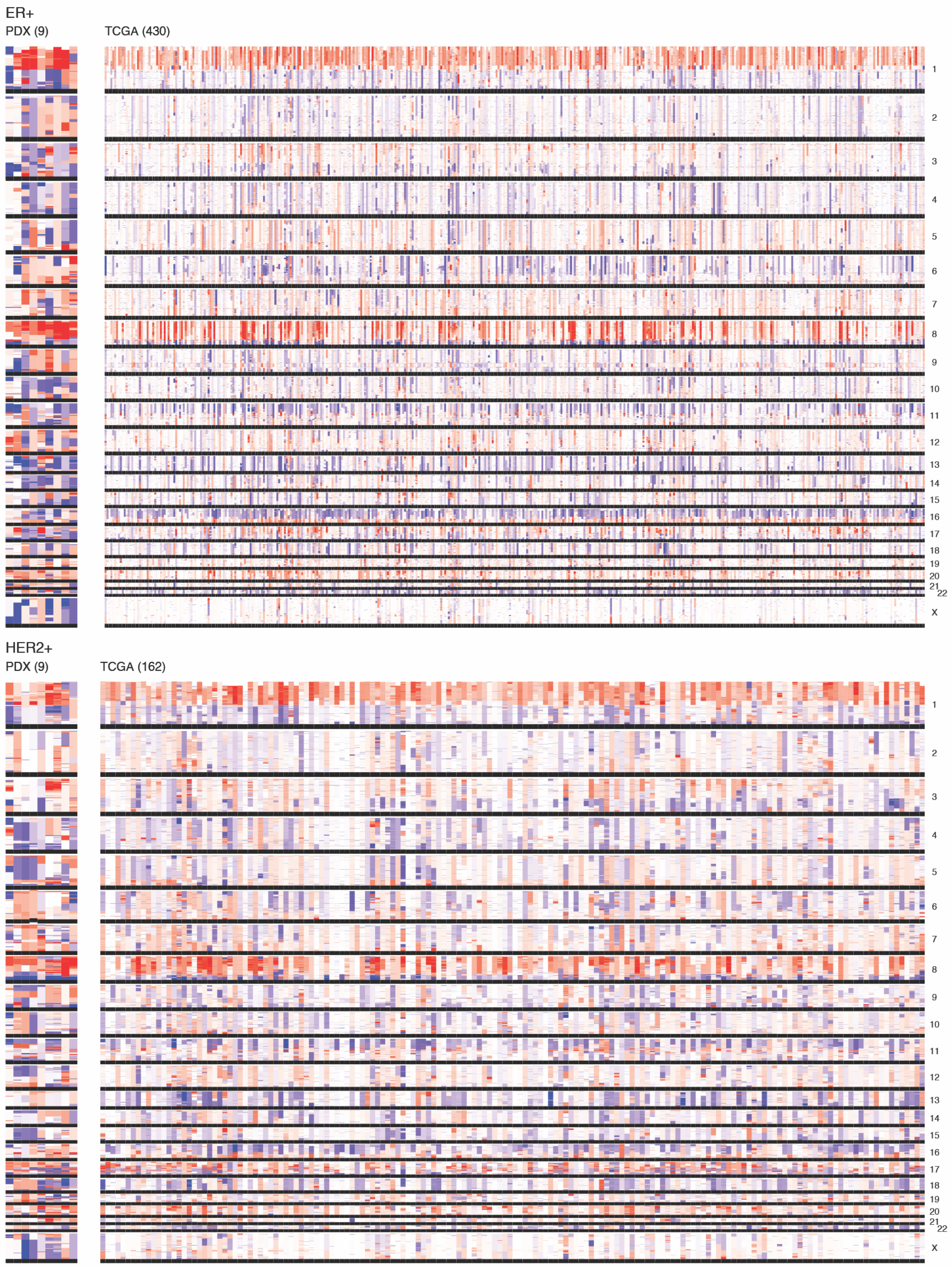

*Supplemental Figure 2:: Mutational Profiles of ER+ (Supplemental Figure 1A) PDXs and TCGA samples and HER2+ (Supplemental Figure 1B) PDX and TCGA samples.*

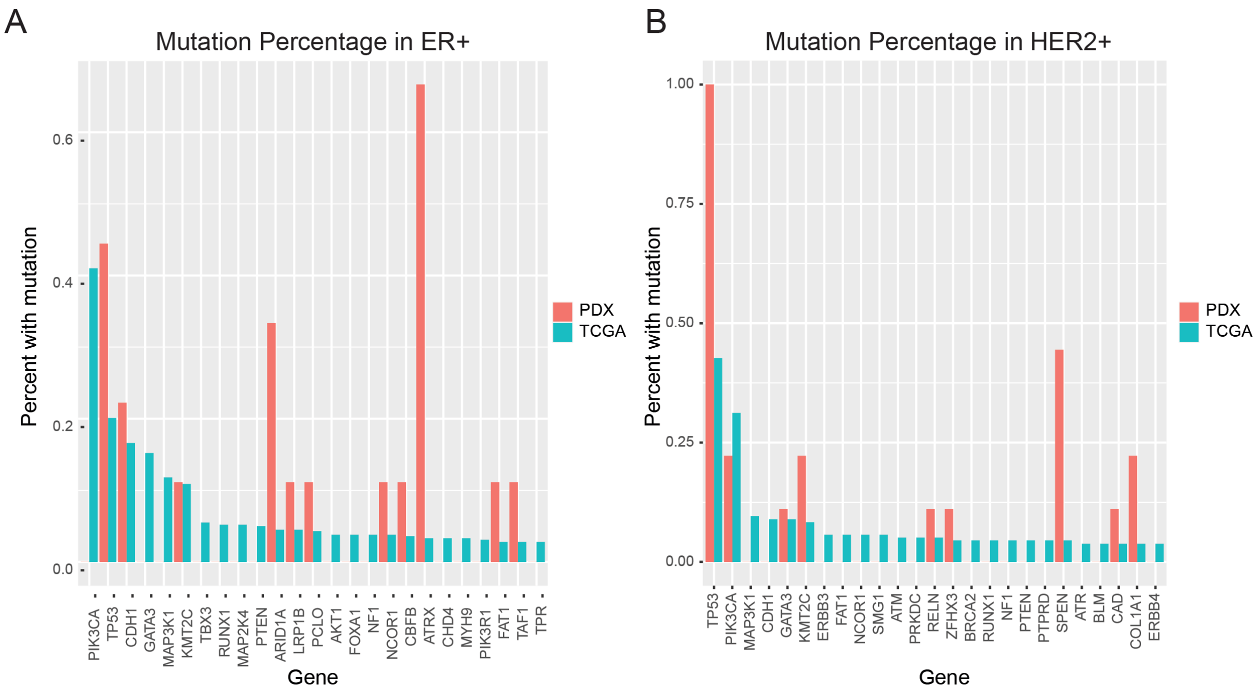

*Supplemental Figure 3: Mutational profiles of ER+ (Supplemental Figure 2A) and HER2+ (Supplemental Figure 2B) PDXs and TCGA samples.*

**Supplemental Table 1**

| Graph | Submodule | CTD connectedness p-value | Linear regression with response | Identified as significant with WGCNA? |
|---|---|---|---|---|
| Carboplatin Epithelial | Submodule 5 | 1.80 e-13 | 0.002 | No |
| Carboplatin Epithelial | Submodule 23 | 8.31 e-11 | 0.014 | Yes |
| Carboplatin Epithelial | Submodule 16 | 8.38 e-21 | 0.015 | Yes |
| Carboplatin Epithelial | Submodule 11 | 3.72 e-18 | 0.029 | No |
| Carboplatin Stromal | Submodule 50 | 4.61 e-05 | 0.005 | No |
| Carboplatin Stromal | Submodule 32 | 3.36 e-05 | 0.036 | No |
| Carboplatin Stromal | Submodule 17 | 1.44 e-12 | 0.047 | No |
| Carboplatin Stromal | Submodule 17 | 3.55 e-05 | 0.049 | No |
| Docetaxel Epithelial | Submodule 23 | 1.53e-18 | 0.010 | No |
| Docetaxel Epithelial | Submodule 52 | 3.54 e-08 | 0.032 | Yes |
| Docetaxel Epithelial | Submodule 32 | 5.75 e-20 | 0.046 | Yes |
| Docetaxel Epithelial | Submodule 24 | 1.59 e-18 | 0.047 | Yes |
| Docetaxel Epithelial | Submodule 43 | 6.77 e-08 | 0.049 | No |
| Docetaxel Stromal | Submodule 49 | 2.03e-10 | 0.0058 | No |
| Docetaxel Stromal | Submodule 17 | 5.66e-07 | 0.024 | Yes |

**Supplemental Table 2**

| Genes | Treatment | Cell Type | Submodule | Informative |
|---|---|---|---|---|
| **F10** | Carboplatin | Epithelial | Submodule 5 | |
| **GLI2** | Carboplatin | Epithelial | Submodule 5 | |
| **DCLK3** | Carboplatin | Epithelial | Submodule 5 | |
| **CERS1** | Carboplatin | Epithelial | Submodule 5 | |
| **C1QL1** | Carboplatin | Epithelial | Submodule 5 | |
| **DENND2A** | Carboplatin | Epithelial | Submodule 5 | |
| **FNDC4** | Carboplatin | Epithelial | Submodule 5 | |
| **MSI1** | Carboplatin | Epithelial | Submodule 5 | X |
| **PLEKHO1** | Carboplatin | Epithelial | Submodule 5 | |
| **TMSB15A** | Carboplatin | Epithelial | Submodule 5 | X |
| **FBN3** | Carboplatin | Epithelial | Submodule 5 | |
| **RNF182** | Carboplatin | Epithelial | Submodule 5 | |
| **DMRT3** | Carboplatin | Epithelial | Submodule 5 | |
| **DMRT2** | Carboplatin | Epithelial | Submodule 5 | |
| **ARHGDIB** | Carboplatin | Epithelial | Submodule 5 | X |
| **LCT** | Carboplatin | Epithelial | Submodule 23 | |
| **LINC02197** | Carboplatin | Epithelial | Submodule 23 | |
| **THRSP** | Carboplatin | Epithelial | Submodule 23 | |
| **ZG16B** | Carboplatin | Epithelial | Submodule 23 | |
| **GGT1** | Carboplatin | Epithelial | Submodule 23 | X |
| **MLLT11** | Carboplatin | Epithelial | Submodule 11 | |
| **SV2A** | Carboplatin | Epithelial | Submodule 11 | x |
| **PYGO1** | Carboplatin | Epithelial | Submodule 11 | |
| **DPY19L2** | Carboplatin | Epithelial | Submodule 11 | |
| **SPSB4** | Carboplatin | Epithelial | Submodule 11 | |
| **ELOVL2** | Carboplatin | Epithelial | Submodule 11 | |
| **NMUR1** | Carboplatin | Epithelial | Submodule 11 | |
| **TAL1** | Carboplatin | Epithelial | Submodule 11 | |
| **TMEFF1** | Carboplatin | Epithelial | Submodule 11 | |
| **GBP5** | Carboplatin | Epithelial | Submodule 16 | |
| **EPHA7** | Carboplatin | Epithelial | Submodule 16 | |
| **FAM218A** | Carboplatin | Epithelial | Submodule 16 | |
| **TDRD12** | Carboplatin | Epithelial | Submodule 16 | |
| **MAGED4** | Carboplatin | Epithelial | Submodule 16 | |

| | | | | |
|---|---|---|---|---|
| **MAGED4B** | Carboplatin | Epithelial | Submodule 16 | |
| **TRO** | Carboplatin | Epithelial | Submodule 16 | |
| **ITPRIPL1** | Carboplatin | Epithelial | Submodule 16 | |
| **PPM1E** | Carboplatin | Epithelial | Submodule 16 | |
| **A430046D13RIK** | Carboplatin | Stromal | Submodule 50 | |
| **LRRC27** | Carboplatin | Stromal | Submodule 50 | |
| **SEC14L2** | Carboplatin | Stromal | Submodule 50 | x |
| **NAPB** | Carboplatin | Stromal | Submodule 62 | |
| **TCEANC** | Carboplatin | Stromal | Submodule 62 | |
| **SERPINI1** | Carboplatin | Stromal | Submodule 32 | x |
| **PLXNA3** | Carboplatin | Stromal | Submodule 32 | |
| **OTUB2** | Carboplatin | Stromal | Submodule 32 | |
| **X4732416N19RIK** | Carboplatin | Stromal | Submodule 17 | |
| **ADAMTS20** | Carboplatin | Stromal | Submodule 17 | x |
| **SLC10A6** | Carboplatin | Stromal | Submodule 17 | |
| **ZFP420** | Carboplatin | Stromal | Submodule 17 | |
| **KLHL4** | Carboplatin | Stromal | Submodule 17 | |
| **PGAP3** | Carboplatin | Stromal | Submodule 17 | |
| **MAOB** | Carboplatin | Stromal | Submodule 17 | |
| **RTN1** | Carboplatin | Stromal | Submodule 17 | |
| **RAI2** | Carboplatin | Stromal | Submodule 17 | |
| **SLC24A3** | Carboplatin | Stromal | Submodule 17 | |
| **CSTAD** | Carboplatin | Stromal | Submodule 17 | |
| **DGKQ** | Carboplatin | Stromal | Submodule 17 | x |
| **MIR99AHG** | Carboplatin | Stromal | Submodule 17 | |
| **GCA** | Carboplatin | Stromal | Submodule 17 | |
| **TM4SF1** | Carboplatin | Stromal | Submodule 17 | |
| **CD300LD3** | Carboplatin | Stromal | Submodule 17 | |
| **E130307A14RIK** | Carboplatin | Stromal | Submodule 26 | x |
| **BC022687** | Carboplatin | Stromal | Submodule 26 | |
| **PODNL1** | Carboplatin | Stromal | Submodule 26 | |
| **ITGA7** | Docetaxel | Epithelial | Submodule 23 | x |
| **METTL7B** | Docetaxel | Epithelial | Submodule 23 | |
| **EFHC2** | Docetaxel | Epithelial | Submodule 23 | |
| **GALNT17** | Docetaxel | Epithelial | Submodule 23 | |
| **LOC100506797** | Docetaxel | Epithelial | Submodule 23 | |
| **GNG4** | Docetaxel | Epithelial | Submodule 23 | |
| **STMND1** | Docetaxel | Epithelial | Submodule 23 | |
| **FBN3** | Docetaxel | Epithelial | Submodule 23 | |

| | | | | |
|---|---|---|---|---|
| **FAM131B** | Docetaxel | Epithelial | Submodule 52 | |
| **SLC10A4** | Docetaxel | Epithelial | Submodule 52 | |
| **KCNE3** | Docetaxel | Epithelial | Submodule 52 | |
| **PDIA2** | Docetaxel | Epithelial | Submodule 52 | |
| **CTAGE6** | Docetaxel | Epithelial | Submodule 32 | |
| **LINC01305** | Docetaxel | Epithelial | Submodule 32 | |
| **C1QL1** | Docetaxel | Epithelial | Submodule 32 | |
| **DCLK3** | Docetaxel | Epithelial | Submodule 32 | |
| **NRG3** | Docetaxel | Epithelial | Submodule 32 | |
| **MCF2L.AS1** | Docetaxel | Epithelial | Submodule 32 | |
| **CERS1** | Docetaxel | Epithelial | Submodule 32 | x |
| **ST8SIA2** | Docetaxel | Epithelial | Submodule 32 | x |
| **IRF4** | Docetaxel | Epithelial | Submodule 32 | |
| **LINC02249** | Docetaxel | Epithelial | Submodule 32 | |
| **ELOVL2** | Docetaxel | Epithelial | Submodule 32 | |
| **NRROS** | Docetaxel | Epithelial | Submodule 24 | |
| **TP53AIP1** | Docetaxel | Epithelial | Submodule 24 | |
| **MAGED4** | Docetaxel | Epithelial | Submodule 24 | x |
| **MAGED4B** | Docetaxel | Epithelial | Submodule 24 | |
| **FAM218A** | Docetaxel | Epithelial | Submodule 24 | |
| **TDRD12** | Docetaxel | Epithelial | Submodule 24 | |
| **EPHA7** | Docetaxel | Epithelial | Submodule 24 | |
| **GBP5** | Docetaxel | Epithelial | Submodule 24 | |
| **SLC26A9** | Docetaxel | Epithelial | Submodule 43 | |
| **SLED1** | Docetaxel | Epithelial | Submodule 43 | |
| **APLN** | Docetaxel | Epithelial | Submodule 43 | |
| **ASB2** | Docetaxel | Epithelial | Submodule 43 | |
| **SLC25A40** | Docetaxel | Stromal | Submodule 49 | |
| **KIF24** | Docetaxel | Stromal | Submodule 49 | x |
| **PARPBP** | Docetaxel | Stromal | Submodule 49 | x |
| **KIF18B** | Docetaxel | Stromal | Submodule 49 | |
| **MCM10** | Docetaxel | Stromal | Submodule 49 | |
| **PAQR8** | Docetaxel | Stromal | Submodule 17 | |
| **KIF24** | Docetaxel | Stromal | Submodule 17 | |
| **PARPBP** | Docetaxel | Stromal | Submodule 17 | |
| **KIF18B** | Docetaxel | Stromal | Submodule 17 | |
| **MCM10** | Docetaxel | Stromal | Submodule 17 | |
| **FOSL1** | Docetaxel | Stromal | Submodule 17 | |
| **HOXC4** | Docetaxel | Stromal | Submodule 17 | |

| | | | |
|---|---|---|---|
| **SIX4** | Docetaxel | Stromal | Submodule 17 |

**Supplemental Table 3**

| PDX Model | Treatment | LogFC Tumor Growth | Response (Modified RECIST) |
|---|---|---|---|
| **BCM-0002** | Untreated | 1.3452 | PD |
| **BCM-0002** | Carboplatin | -0.5891 | PR |
| **BCM-0002** | Docetaxel20 | -0.1008 | SD |
| **BCM-0104** | Untreated | 2.4278 | PD |
| **BCM-0104** | Carboplatin | -11.7735 | CR |
| **BCM-0104** | Docetaxel20 | -5.0307 | PR |
| **BCM-0132** | Untreated | 2.3263 | PD |
| **BCM-0132** | Carboplatin | 1.5665 | PD |
| **BCM-0132** | Docetaxel20 | -0.8381 | PR |
| **BCM-2147** | Untreated | 3.2497 | PD |
| **BCM-2147** | Carboplatin | -11.2993 | CR |
| **BCM-2147** | Docetaxel20 | 1.4375 | PD |
| **BCM-2665** | Untreated | 2.3624 | PD |
| **BCM-2665** | Carboplatin | 1.9106 | PD |
| **BCM-2665** | Docetaxel20 | -0.2426 | SD |
| **BCM-3107** | Untreated | 1.0812 | PD |
| **BCM-3107** | Carboplatin | -3.3873 | PR |
| **BCM-3107** | Docetaxel20 | -6.6821 | PR |
| **BCM-3204** | Untreated | 1.1731 | PD |
| **BCM-3204** | Carboplatin | -1.2278 | PR |
| **BCM-3204** | Docetaxel20 | -1.0264 | PR |
| **BCM-3469** | Untreated | 2.0463 | PD |
| **BCM-3469** | Carboplatin | 0.5562 | PD |
| **BCM-3469** | Docetaxel20 | -1.1147 | PR |
| **BCM-3561** | Untreated | 2.0856 | PD |
| **BCM-3561** | Carboplatin | 0.2887 | PD |
| **BCM-3561** | Docetaxel20 | -4.4062 | PR |
| **BCM-3611** | Untreated | 1.9198 | PD |
| **BCM-3611** | Carboplatin | 1.1086 | PD |
| **BCM-3611** | Docetaxel20 | -5.2165 | PR |
| **BCM-3887** | Untreated | 2.8573 | PD |
| **BCM-3887** | Carboplatin | -6.5797 | PR |

| | | | |
|---|---|---|---|
| **BCM-3887** | Docetaxel20 | -8.5532 | PR |
| **BCM-3904** | Untreated | 2.2578 | PD |
| **BCM-3904** | Carboplatin | 0.4213 | PD |
| **BCM-3904** | Docetaxel20 | -11.0123 | CR |
| **BCM-3936** | Untreated | 2.169 | PD |
| **BCM-3936** | Carboplatin | 1.6777 | PD |
| **BCM-3936** | Docetaxel20 | -1.1577 | PR |
| **BCM-4013** | Untreated | 1.4387 | PD |
| **BCM-4013** | Carboplatin | -3.4795 | PR |
| **BCM-4013** | Docetaxel20 | -6.5906 | PR |
| **BCM-4195** | Untreated | 2.7993 | PD |
| **BCM-4195** | Carboplatin | -11.563 | CR |
| **BCM-4195** | Docetaxel20 | -2.8592 | PR |
| **BCM-4272** | Untreated | 2.0187 | PD |
| **BCM-4272** | Carboplatin | -1.2922 | PR |
| **BCM-4272** | Docetaxel20 | -3.6364 | PR |
| **BCM-4664** | Untreated | 2.659 | PD |
| **BCM-4664** | Carboplatin | 1.9476 | PD |
| **BCM-4664** | Docetaxel20 | -1.9567 | PR |
| **BCM-4913** | Untreated | 2.0279 | PD |
| **BCM-4913** | Carboplatin | -11.6772 | CR |
| **BCM-4913** | Docetaxel20 | -9.4681 | PR |
| **BCM-5471** | Untreated | 2.027 | PD |
| **BCM-5471** | Carboplatin | 1.744 | PD |
| **BCM-5471** | Docetaxel20 | 1.7511 | PD |
| **BCM-5998** | Untreated | 1.4783 | PD |
| **BCM-5998** | Carboplatin | 1.1104 | PD |
| **BCM-5998** | Docetaxel20 | 0.1179 | SD |
| **BCM-7482** | Untreated | 2.3761 | PD |
| **BCM-7482** | Carboplatin | -4.7806 | PR |
| **BCM-7482** | Docetaxel20 | 1.8819 | PD |
| **BCM-7649** | Untreated | 2.5024 | PD |
| **BCM-7649** | Carboplatin | 2.0707 | PD |
| **BCM-7649** | Docetaxel20 | 0.945 | PD |
| **BCM-7821** | Untreated | 1.5545 | PD |
| **BCM-7821** | Carboplatin | 1.0228 | PD |
| **BCM-7821** | Docetaxel20 | 0.3032 | PD |
| **BCM-9161** | Untreated | 2.0945 | PD |
| **BCM-9161** | Carboplatin | -11.804 | CR |

| | | | |
|---|---|---|---|
| **BCM-9161** | Docetaxel20 | -9.595 | PR |
| **BCM-15003** | Untreated | 0.4757 | PD |
| **BCM-15003** | Carboplatin | -0.72 | PR |
| **BCM-15003** | Docetaxel20 | -5.1363 | PR |
| **BCM-15006** | Untreated | 1.1334 | PD |
| **BCM-15006** | Carboplatin | -3.7263 | PR |
| **BCM-15006** | Docetaxel20 | -2.3104 | PR |
| **BCM-15008** | Untreated | 1.6459 | PD |
| **BCM-15008** | Carboplatin | -7.3592 | PR |
| **BCM-15008** | Docetaxel20 | -5.7082 | PR |
| **BCM-15029** | Untreated | 0.7053 | PD |
| **BCM-15029** | Carboplatin | -0.3449 | SD |
| **BCM-15029** | Docetaxel20 | -0.8016 | PR |
| **HCI-001** | Untreated | 2.6101 | PD |
| **HCI-001** | Carboplatin | -11.5077 | CR |
| **HCI-001** | Docetaxel20 | 0.9191 | PD |
| **HCI-002** | Untreated | 2.2693 | PD |
| **HCI-002** | Carboplatin | 1.3226 | PD |
| **HCI-002** | Docetaxel20 | -5.5394 | PR |
| **HCI-009** | Untreated | 0.6546 | PD |
| **HCI-009** | Carboplatin | 0.9697 | PD |
| **HCI-009** | Docetaxel20 | -11.7782 | CR |
| **HCI-010** | Untreated | 1.4172 | PD |
| **HCI-010** | Carboplatin | -0.1587 | SD |
| **HCI-010** | Docetaxel20 | -1.5732 | PR |
| **HCI-015** | Untreated | 1.4679 | PD |
| **HCI-015** | Carboplatin | -11.6033 | CR |
| **HCI-015** | Docetaxel20 | -1.9634 | PR |
| **HCI-016** | Untreated | 1.8509 | PD |
| **HCI-016** | Carboplatin | -2.8261 | PR |
| **HCI-016** | Docetaxel20 | -9.5717 | PR |
| **HCI-019** | Untreated | 1.0277 | PD |
| **HCI-019** | Carboplatin | -2.6227 | PR |
| **HCI-019** | Docetaxel20 | -4.6395 | PR |
| **HCI-023** | Untreated | 1.1728 | PD |
| **HCI-023** | Carboplatin | -1.7487 | PR |
| **HCI-023** | Docetaxel20 | -3.2258 | PR |
| **HCI-024** | Untreated | 0.7037 | PD |
| **HCI-024** | Carboplatin | -0.0272 | SD |

| | | | |
|---|---|---|---|
| **HCI-024** | Docetaxel20 | -6.3931 | PR |
| **HCI-027** | Untreated | 2.6661 | PD |
| **HCI-027** | Carboplatin | 1.8783 | PD |
| **HCI-027** | Docetaxel20 | -11.7357 | CR |
| **HCI-028** | Untreated | 1.9808 | PD |
| **HCI-028** | Carboplatin | 1.3713 | PD |
| **HCI-028** | Docetaxel20 | 1.5567 | PD |
| **HCI-030** | Untreated | 1.0243 | PD |
| **HCI-030** | Carboplatin | -11.7073 | CR |
| **HCI-030** | Docetaxel20 | -11.6133 | CR |
| **WHIM2** | Untreated | 2.5855 | PD |
| **WHIM2** | Carboplatin | 2.3627 | PD |
| **WHIM2** | Docetaxel20 | -3.002 | PR |
| **WHIM6** | Untreated | 1.8975 | PD |
| **WHIM6** | Carboplatin | -11.2404 | CR |
| **WHIM6** | Docetaxel20 | 0.8576 | PD |
| **WHIM14** | Untreated | 1.0744 | PD |
| **WHIM14** | Carboplatin | -8.1665 | PR |
| **WHIM14** | Docetaxel20 | -6.3467 | PR |
| **WHIM30** | Untreated | 2.1474 | PD |
| **WHIM30** | Carboplatin | -10.2041 | CR |
| **WHIM30** | Docetaxel20 | -9.8985 | PR |
| **WHIM68** | Untreated | 1.4674 | PD |
| **WHIM68** | Carboplatin | -11.5574 | CR |
| **WHIM68** | Docetaxel20 | -5.6538 | PR |
